# Pioneer activity distinguishes activating from non-activating pluripotency transcription factor binding sites

**DOI:** 10.1101/2022.07.27.501606

**Authors:** Michela Maresca, Teun van den Brand, Hangpeng Li, Hans Teunissen, James Davies, Elzo de Wit

**Affiliations:** Division of Gene Regulation, Oncode Institute & The Netherlands Cancer Institute, Amsterdam, the Netherlands; MRC Molecular Haematology Unit, MRC Weatherall Institute of Molecular Medicine, Radcliffe Department of Medicine, University of Oxford, Oxford, UK

**Author notes:** these authors contributed equally (Michela Maresca, Teun van den Brand).

## Abstract

Genome-wide transcriptional activity involves the binding of many transcription factors to thousands of sites in the genome. Determining which sites are directly driving transcription remains a challenge. Here we use acute protein depletion of the pioneer transcription factors OCT4 and SOX2 to establish their functionality in maintaining chromatin accessibility. We show that thousands of accessible sites are lost within an hour of protein depletion, indicating rapid turnover of these sites in the absence of pioneer factors. To understand the relationship with transcription we performed nascent transcription analysis and found that open chromatin sites that are maintained by SOX2 are highly predictive of gene expression, in contrast to SOX2 binding sites that do not maintain accessibility. We use CRISPR-Cas9 genome editing in the *Klf2* locus to functionally validate a predicted regulatory element. We conclude that the regulatory activity of SOX2 is exerted largely at sites where it maintains accessibility and that other binding sites are largely dispensable for gene regulation.

## Introduction

In a given cell, gene expression is regulated by the concerted action of hundreds of transcription factors (TFs)(Holmberg and Perlmann, 2012). Sequence-specific TFs enhance transcription by directly binding to DNA and promoting the recruitment of the transcriptional machinery(Allen and Taatjes, 2015; Grunstein, 1997; Isbel et al., 2022; Ptashne and Gann, 1997). Mammalian genomes encode over a thousand different TFs (Vaquerizas et al., 2009) and cell type specific TFs to a large degree determine cellular states. Therefore, delineating cell type specific activity of TFs is crucial to understanding organismal development and homeostasis. Discriminating functional TF binding sites, i.e. those that activate transcription, from non-functional sites that do not activate transcription, is a key challenge of the post-genomic era.

Classically, TF binding specificity has been characterized with different complementary methods. Proteins DNA interactions can be measured *in vitro* with methods such as SELEX or protein binding microarrays (Stormo and Zhao, 2010). Alternatively, one can determine the binding sites of DNA binding factors in cellular context using chromatin immunoprecipitation (ChIP), which, when coupled to genome-wide read-outs, (Blat and Kleckner, 1999; Ren et al., 2000) enables their genome-wide identification. For the majority of transcription factors, this results in the identification of tens of thousands of bound sites. For most transcription factors, the number of bound sites is substantially lower than the number of DNA sequences in the genome that correspond to the *in vitro* defined binding motif, indicating that these factors are not recruited to all motifs with equal efficiency. Furthermore, the same sequence-specific transcription factor can have different binding profiles depending on the cell type (Lodato et al., 2013). Moreover, it should be noted that the number of regulatory elements that control gene expression (i.e. functional elements) is again smaller than the number of bound sites (Todd et al., 2019).

One of the reasons why not all motifs for a given transcription factor are bound, is that the DNA is not accessible for binding by the factor in question. To make DNA elements accessible, a special class of transcription factors called pioneer factors are required (Zaret and Carroll, 2011; Zaret and Mango, 2016). Pioneer factors can bind to DNA wrapped around the histone octamer. Upon binding, these factors recruit chromatin remodeling complexes (Swinstead et al., 2016) resulting in the formation of nucleosome free, accessible, DNA. These genomic regions can then be bound by non-pioneer transcription factors to activate gene expression. During differentiation, the transition from one cell type to another is driven by the formation of cell type specific accessible regions, which, in turn, activate cell type specific genes. As such, activity of pioneer factors often represents the first step in cell fate commitment (Zaret and Mango, 2016). Reprogramming of somatic cells to induced pluripotent stem cells is also preceded by the induction of pluripotency associated accessible DNA elements (Li et al., 2017, 2021). It is therefore not surprising that three out of four of the original Yamanaka reprogramming factors are pioneer transcription factors (i.e. OCT4, SOX2 and KLF4) (Takahashi and Yamanaka, 2006).

OCT4 and SOX2, together with NANOG are considered the master regulators of pluripotency (Boyer et al., 2005; Kim et al., 2008). They cooperate to maintain transcription of pluripotency genes and repress expression of developmental genes (Chambers et al., 2003; Loh et al., 2006; Mitsui et al., 2003; Niwa et al., 2000; Schöler et al., 1990). During differentiation, the genes encoding these factors are downregulated and loss-of-function studies show that mouse embryonic stem cells (mESCs) depend on the expression of OCT4 and SOX2 to maintain the pluripotent state (Avilion et al., 2003; Nichols et al., 1998). Genome-wide occupancy analysis by ChIPseq revealed binding of OCT4 and SOX2 close to other pluripotency-associated genes (Marson et al., 2008). CryoEM structures of the DNA binding domains of the OCT4-SOX2 dimer binding to nucleosomes show that their binding facilitates the local distortion of the nucleosomal DNA (Dodonova et al., 2020; Michael et al., 2020), indicating the initial steps towards generating open chromatin sites. Knock-out of OCT4 leads to a loss of accessible sites (King and Klose, 2017), further emphasizing their role as pioneer factors.

Accessible chromatin sites, which are created by pioneer factors during development and can be measured with ATACseq or DNAse I hypersensitivity mapping, clearly correlate with gene expression changes at selected loci (see for example: (Isoda et al., 2017; Pijuan-Sala et al., 2020)). However, identifying the exact sites that drive the expression changes remains difficult. For instance, experiments that aimed to link changes in the open chromatin landscape to changes in steady state RNA levels in a breast cancer cell line following treatment with retinoic acid or TFG-beta for 72 hours found only a meagre correlation (Kiani et al., 2022). Loss-of-function studies for transcription factors suffer from a related complication. During conditional knock-out or knock-down experiments the protein level is passively decreased through cell division or the rate of degradation, which means it can take many hours before the protein is completely depleted. Because loss of pluripotency TFs induces differentiation, this can result in heterogeneity between cells, making it difficult to separate direct from indirect effects. In recent years, acute protein depletion methods have been developed which can ablate proteins directly (Verma et al., 2020). These typically work by using a small molecule to target the protein of interest to an E3 ligase complex resulting in ubiquitination and proteasomal degradation. This results in rapid, synchronous and near complete loss of a protein.

In this work we used acute transcription factor depletion to determine the dynamics of open chromatin sites created by pioneer factors and how this is related to gene expression. Acute loss of OCT4, SOX2 and NANOG in mouse embryonic stem cells (mESCs) reveals that thousands of accessible sites are lost in less than two hours upon depletion of OCT4 and SOX2. When we link the changes in open chromatin following SOX2 depletion to changes in nascent transcription we find that differentially accessible chromatin sites are strongly predictive for changes in gene expression. Importantly, binding sites of SOX2 that are not associated with changes in accessibility have poor predictive power for changes in gene expression. Our experiments clearly establish the importance of chromatin accessibility created and continuously maintained by pioneer transcription factors in the regulation of gene expression.

## Results

### Continuous pioneering activity of transcription factors is necessary for maintenance of accessible chromatin

To determine the direct consequences of transcription factor (TF) loss at high temporal resolution we employed the degradation tag (dTAG) system to induce rapid and acute protein depletion (Nabet et al., 2018). We previously performed C-terminal tagging of the *Sox2* and *Nanog* genes with the *FKBP^F36V^* sequence using CRISPR/Cas9 genome editing in mouse embryonic stem cells (mESC) (Liu et al., 2020). The heterobifunctional molecule dTAG-13 binds to the FKBP^F36V^ domain and targets the fusion protein for proteasomal degradation (Figure 1A). In addition to the SOX2 and NANOG degron lines that we created we also included a previously published OCT4 degron line (Boija et al., 2018) in our analyses. Immunoblotting shows that treatment with dTAG efficiently degrades tagged protein levels within one hour (Figure 1B). Although SOX2 and OCT4 become undetectable on Western, for NANOG we still detect some residual protein expression (Figure 1B). To determine whether dTAG treatment is uniquely selective for the fusion proteins we performed quantitative mass spectrometry-based proteomics after 30 min of protein depletion (Figure 1C, Figure S1A). For the SOX2 line, after 30 minutes of dTAG treatment, the only protein that showed significant changes was SOX2. For the OCT4 and NANOG depletion, we find that there are additional proteins whose level is affected after 30 minutes of dTAG treatment. However, none of these proteins are shared between the two lines, suggesting that these are not caused by off-target degradation induced by the dTAG small molecule treatment, but rather a secondary effect specific to the depleted protein. Collectively, these results show that acute protein depletion of pluripotency transcription factors is fast and highly specific.

**Figure 1.**
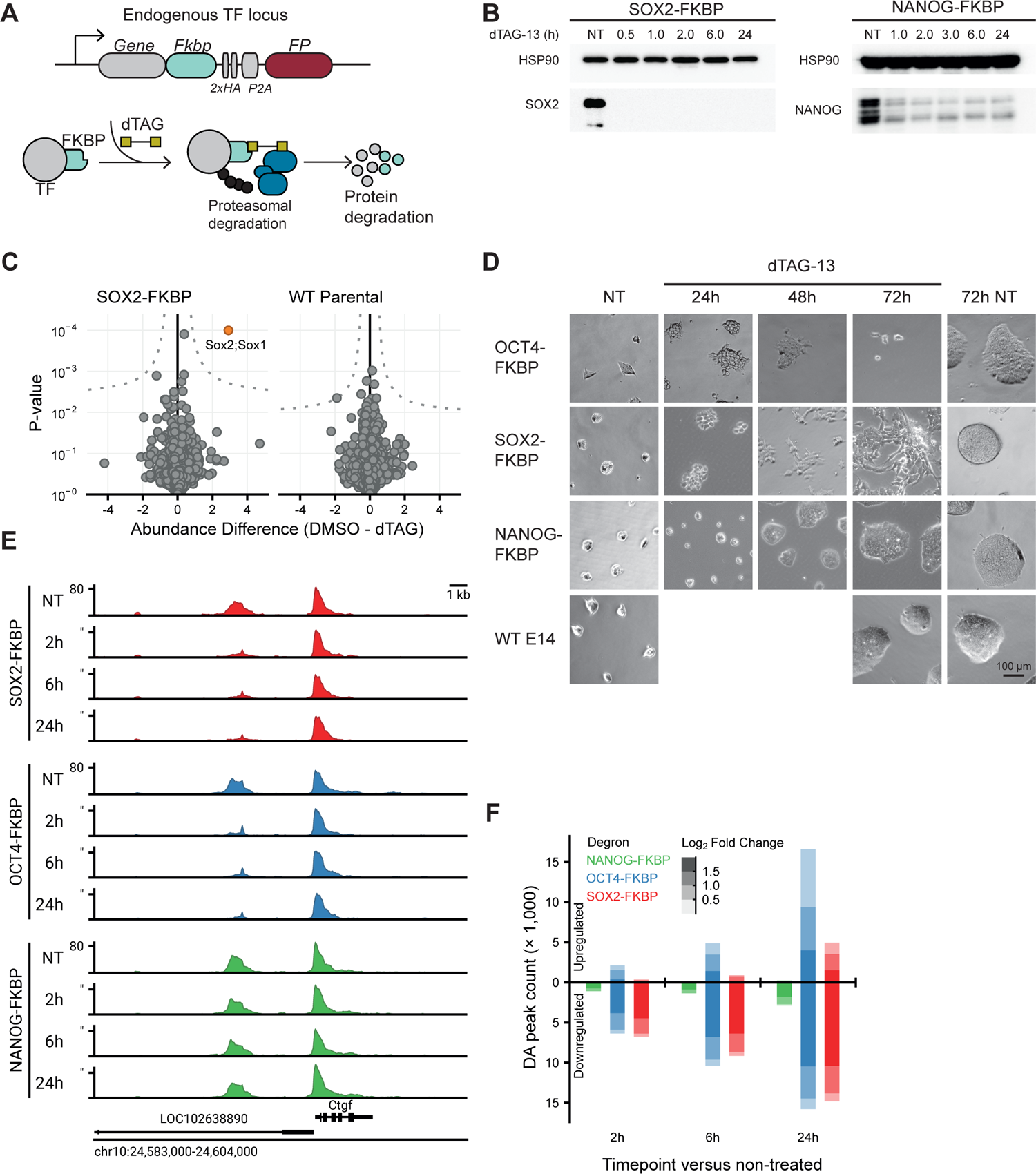
Rapid depletion of SOX2 and OCT4 affects the accessibility landscape of thousands of sites. **(A)** Schematic representation of the dTAG system, wherein an FKBP-tagged protein can be proteasomally degraded upon addition of the dTAG-13 small molecule. **(B)** Western blot showing protein abundance of SOX2 and NANOG upon addition of dTAG-13 for the indicated times in respective FKBP-tagged cell lines. HSP90 was used as loading control. NT = Not Treated. **(C)** Quantitative mass spectrometry results showing the differential protein abundance upon 30 minutes of dTAG treatment versus DMSO treated SOX2-FKBP cells and parental (untagged) cells. **(D)** Brightfield images showing colony morphology upon addition of dTAG-13 for the indicated timepoints. **(E)** Genomic tracks showing accessibility by ATACseq around the *Ctgf* gene for FKBP-tagged cell lines after indicated times of depletion by dTAG treatment. Y-axes show reads per genomic content (RPGC) **(F)** The number of differentially accessible peaks shown for every FKBP-tagged cell line and time of depletion. Positive bars indicate peaks with increased accessibility (gain), negative bars peaks with decreased accessibility (loss). Colour intensity is indicative of effect size of change in accessibility.

To understand the cell biological consequences of pluripotency TF depletion we determined with brightfield microscopy whether there were any morphological changes (Figure 1D). While mESCs typically grow in round colonies, in the absence of SOX2 or OCT4 the ES colonies presented a more rugged morphology, indicating on-going differentiation (Avilion et al., 2003; Keramari et al., 2010). At 72 hours post depletion (hpd) of OCT4, hardly any cells could be observed in culture, showing that prolonged and complete loss of OCT4 is detrimental to these cells. In contrast, SOX2 loss did not lead to cell death after 72 hours of dTAG treatment, but did show an almost complete loss of colony structure. Compared to OCT4 and SOX2, NANOG depletion had no obvious impact on ES cell colony morphology (Figure 1D). These observations reveal a clear difference in consequences for colony morphology and cellular growth between the pioneer TFs OCT4 and SOX2 on the one hand and NANOG on the other.

To determine the genome-wide molecular consequences of loss of these three TFs we performed ATACseq. This allowed us to profile the direct effects that each TF has on the accessible chromatin of mESCs. In ATACseq the bacterial transposase Tn5 tagments DNA that is accessible enabling the detection of open chromatin regions (OCRs). We performed an ATACseq time course experiment at 2, 6 and 24 hours following OCT4, SOX2 or NANOG depletion. The genomic region surrounding the *Ctgf* locus is exemplary for the consequences of acute TF loss on the chromatin accessibility landscape (Figure 1E). An OCR ∼4.5 kb upstream from the *Ctgf* TSS starts losing accessibility already after 2 hours of SOX2 or OCT4 depletion, but is unaffected by NANOG depletion. To catalogue the genome-wide changes upon TF depletion we perform peak calling using MACS2, which detected 184,964 peaks on the combined time series of the three degron lines. After filtering, we called differentially accessible regions (DARs) on 166,267 peaks using DESeq2 (see Methods for details). Figure 1F shows the distribution of DARs over time for the different TF degrons. Depletion of the non-pioneer factor NANOG affects accessible chromatin at a limited number of regions over the investigated time course (n=3,118 at 24hpd). On the other hand, depletion of OCT4 and SOX2 results in loss of accessibility of thousands of regions already at 2hpd (n=6,757 and n=6,362, respectively), in line with their role as pioneer factors. The dichotomy in pioneer activity between OCT4 and SOX2 on the one hand and NANOG on the other is also seen in quantitative analyses (Figure S1B, S1C). Over time, depletion of OCT4 and SOX2 also results in the establishment of 16,620 and 4,963 newly accessible sites at 24hpd, respectively (Figure 1F). Because the increases in accessibility happen mostly at the later timepoints this is likely a secondary effect of SOX2 and OCT4 depletion, with the primary consequence being loss of accessibility. Furthermore, the surprisingly fast loss of accessible chromatin in the absence of pioneer TFs suggests that continuous pioneer activity is required for the maintenance of these OCRs.

### SOX2 and OCT4 can maintain accessibility cooperatively and independently

In pluripotent cells, OCT4 and SOX2 often occupy the same genomic sites (Chen et al., 2008; Marson et al., 2008). Furthermore, Cryo-EM structures of the DNA binding domains in complex with nucleosomes have revealed that OCT4 and SOX2 form a heterodimer that distorts the interface between DNA and the nucleosome, which may promote the formation of an accessible site (Dodonova et al., 2020; Michael et al., 2020). The rapid and specific depletion of OCT4 and SOX2 enables us to determine the contribution of either factor to the formation of OCRs in a cellular context. To this end, we investigated the DARs at 2hpd in more detail and stratified them into different categories. Most prominent are the sites that lose accessibility following depletion of either OCT4 or SOX2 and thus depend on the action of both proteins (Figure. 2A). On the other hand, we observed a sizable number of peaks that are exclusively lost following depletion of SOX2 or OCT4, but not in both, making these uniquely dependent on either factor (Figure. 2A). To identify different types of OCR we used statistical interaction effects to define DARs that depend exclusively on SOX2 or OCT4, or that depend on both for maintaining accessibility (Figure 2B). The majority of DARs (n=2,852) rely on both OCT4 and SOX2 to maintain an open chromatin state (DAR^OCT4/SOX2^), 1,431 sites rely on SOX2 alone to establish open chromatin (DAR^SOX2^), while a small number of regions (n=343) depend only on OCT4 (DAR^OCT4^). It should be noted that we apply a stringent, buffered cut-off for these specific categories, which means our list is likely non-exhaustive, but minimizes ambiguity between the categories.

**Figure 2.**
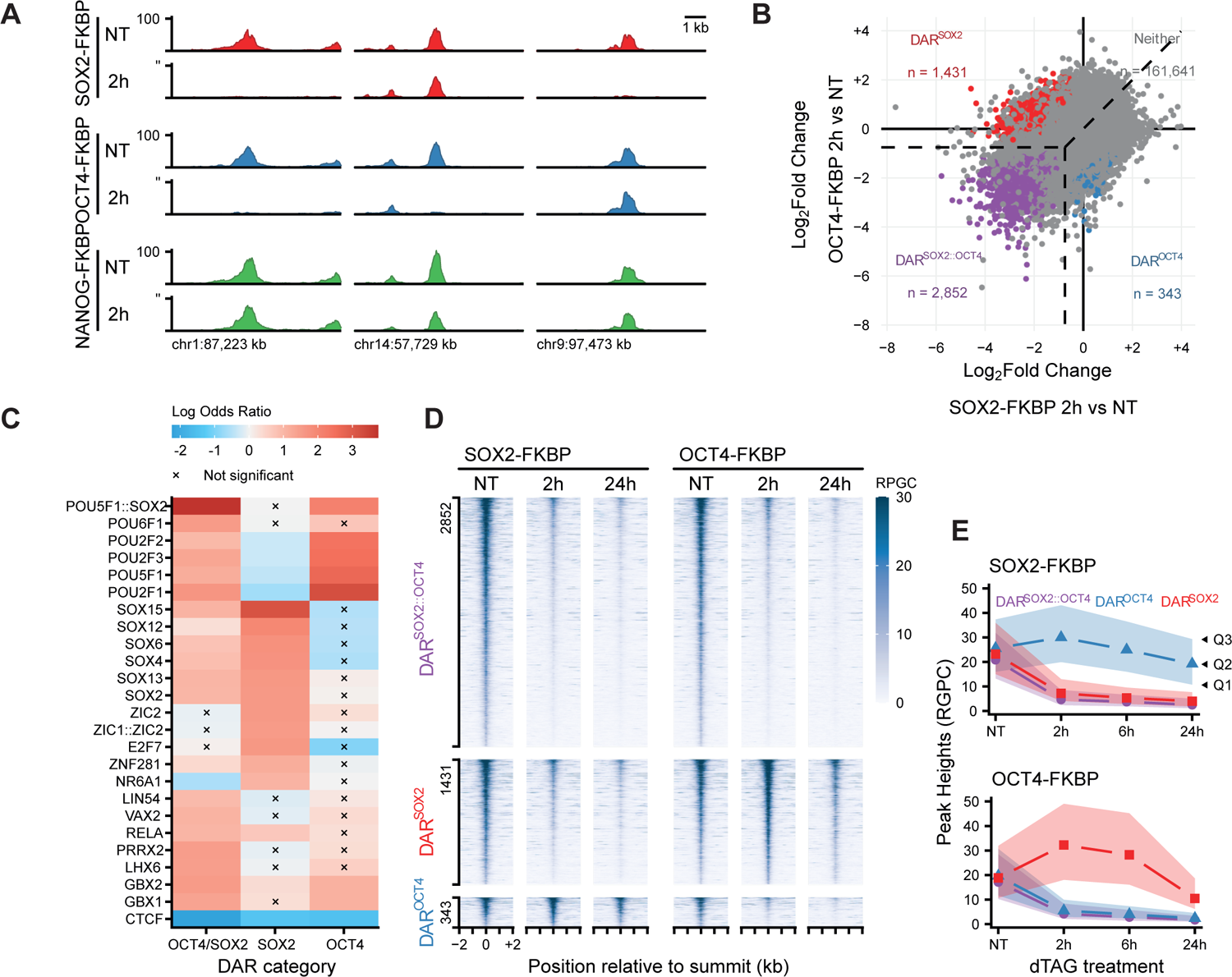
SOX2 and OCT4 regulate accessibility both cooperatively and independently. **(A)** Genomic tracks showing accessibility by ATACseq in untreated cells and after 2 hours of depletion in for SOX2-, OCT4- and NANOG-FKBP-tagged cell lines. Examples of sites where accessibility is regulated by both, or exclusively, SOX2 or OCT4 are shown. Y-axes show reads per genomic content (RPGC) **(B)** Scatter plot showing the effect sizes at peak locations following 2 hours of SOX2 depletion against 2 hours of OCT4 depletion. Colours indicate categorisation of differentially accessible regions (DARs) as mentioned in the main text. **(C)** Heatmap showing the enrichment and depletion of TF binding motifs in the DARs. Shown are the top 25 motifs with the lowest enrichment p-value in any category. Log odds ratio was calculated as coefficients in logistic regression with DAR category membership as dependent variable, controlling for the GC-bias and total motif presence in peaks. Significance was tested using a single-parameter Wald-test in the same regression. **(D)** Tornado plots showing accessibility at peaks belonging to the DAR categories in the SOX2 and OCT4 degron cell lines. **(E)** Quantification of the heights of peaks belonging to the DAR categories over time of depletion in the SOX2 and OCT4 degron lines. Shaded area indicates the first quartile (Q1) up to the third quartile (Q3) of heights, whereas points indicate the median (Q2).

TFs identify their target sites by recognition of a specific DNA sequence motif. To determine whether the three different categories of DARs that we identified, are a consequence of a difference in sequence context, we performed motif enrichment analysis (Figure 2C). The DAR^OCT4/SOX2^ sites are strongly enriched for the OCT4::SOX2 compound motif, consistent with the model that, at those sites, the two proteins are co-localizing and acting in heterodimerised form (Michael et al., 2020).The DAR^SOX2^ category instead shows a stronger enrichment for motifs of SOX family members compared to the combined motif, with the strongest enrichment for SOX15, which carries the canonical ACAAT/ATTGT sequence. The DAR^OCT4^, on the other hand, are more strongly enriched for motifs of POU family members (to which OCT4 belongs). SOX2 and OCT4 may act as monomers or homodimers in establishing chromatin accessibility at the DAR^SOX2^ and DAR^OCT4^ sites, respectively. Alternatively, they could form within-family heterodimers, which would be difficult to detect given the similar binding sequence. Besides SOX and OCT motifs we also find enrichment for additional motifs (Figure 2C). These TFs may cooperate with SOX2 or OCT4 to maintain open chromatin or could be secondary consequences of changes in the level or activity of these factors.

The stratification of DARs enables us to track their dynamics over time following TF depletion. As expected, the DAR^OCT4/SOX2^ sites show a rapid and long-term decrease in accessibility following either SOX2 or OCT4 depletion (Figure 2D). We can now also track what happens to the sites that are likely regulated by either TF alone. Intriguingly, following 2 and 6 hours of OCT4 depletion we find that the DAR^SOX2^ are increased compared to the untreated samples (Figure 2D,E). However, at 24hpd the accessibility of DAR^SOX2^ sites is decreased 3-fold compared to the levels at 2hpd, but still higher compared to DAR^OCT4/SOX2^. In a similar fashion we determined the accessibility dynamics of the DAR^OCT4^ sites following SOX2 depletion. Although, the trend of the DAR^OCT4^ sites is similar (i.e. an increase followed by a decrease) the magnitude of the effect is much smaller (1.5-fold) compared to the DAR^SOX2^ sites following OCT4 depletion (Figure 2E). To better understand the differences in accessibility dynamics between the different lines we determined the levels of the OCT4 and SOX2 protein following the depletion of these factors. We found that following the depletion of OCT4, the level of SOX2 protein is not affected (Figure S2). Also, when SOX2 is depleted, the level of OCT4 remains stable in the period we assayed (first 24hpd). This may explain why the independent sites of the non-depleted factor remain open compared to cooperative sites and the independent sites of the depleted factor. The temporary increase in accessibility for DAR^SOX2^ sites could mean that following the depletion of OCT4, SOX2 is freed up and redistributed to sites where they do not bind together, leading to a temporary increase in accessibility. Taken together, we show that acute synchronized depletion can identify complex relationships that are obscured in depletions over longer time scales.

### SOX2 directly controls accessibility with a sub-hour time resolution

Because our analyses indicated that depletion of SOX2 maintains accessibility together with OCT4 but also at a large set of sites independent of OCT4 we decided to further explore the role of SOX2 in transcription regulation. The sites that lose accessibility following 2 hours of SOX2 depletion, number in the thousands (Figure 1F), suggesting that at this time point we have already identified the vast majority of sites whose accessibility is controlled by SOX2. Therefore, we decided to increase our temporal resolution for SOX2 depletion by analyzing two additional time points before 2 hours, at 30 minutes and 1 hour from the same time series. Visual inspection of the data already revealed that there are different temporal dynamics between sites with respect to their response following SOX2 depletion (Figure 3A). For a subset of sites, no residual accessibility can be detected as soon as after 30 minutes of depletion, whereas other sites take more than 2 hours to become fully inaccessible. Quantification of the DARs at every time point shows a gradual loss of OCRs over time (Figure 3B). The gain of accessibility in OCRs following SOX2 depletion is very limited (less than 500 sites in the first 2hpd), and only at 24hpd thousands of OCR increase their accessibility.

**Figure 3.**
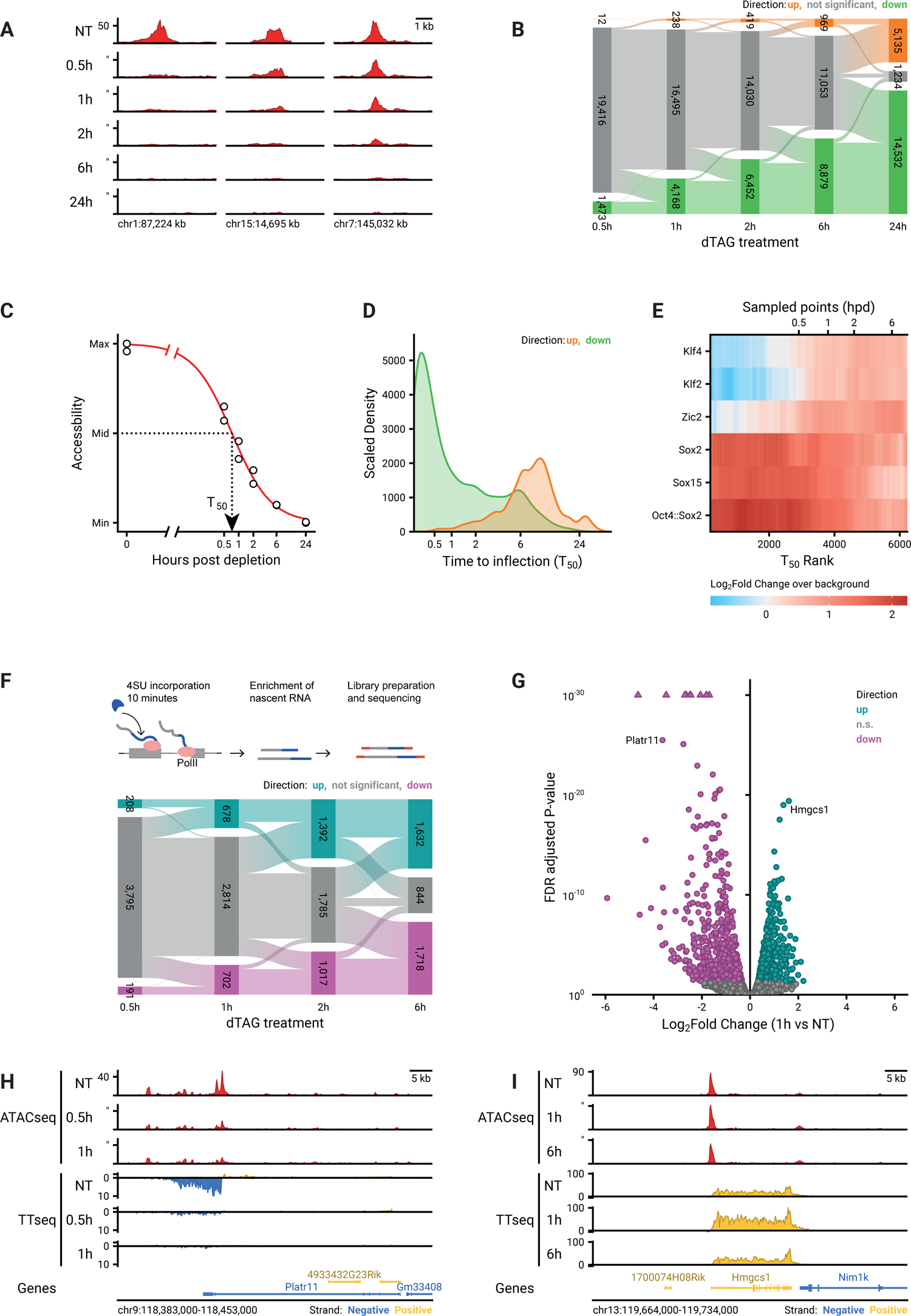
Loss of SOX2 affect chromatin accessibility and transcription with sub-hour time resolution. **(A)** Example regions of changes in accessibility as measured by ATACseq during the SOX2 depletion time course. Y-axis indicates reads per genomic content (RGPC). **(B)** Alluvial diagram shows the number of regions gaining accessibility (up, orange) or losing accessibility (down, green) over the SOX2 depletion time course. **(C)** Illustration of estimating the time to inflection (T_50_) for a DAR, wherein a logistic model is fit to the data over log-time after depletion. **(D)** Kernel density estimate of the time to inflection for DARs going up and down. The density is scaled such that the area under the curve integrates to the number of DARs. Time to inflection is displayed with inverse hyperbolic sine transformation. **(E)** Heatmap of motif enrichment displaying downregulated DARs on the x-axis sorted by time to inflection. Colours display a fold change of a centered running mean, measuring the fraction of peaks containing the indicated motifs over a 400 peaks window, relative to the fraction of nDARs containing these motifs. Sampled time points are indicated at the bins where the time to inflection surpasses the sampled time. **(F)** Top: Schematic representation of the TT_chem_seq method to measure nascent transcripts. Bottom: Alluvial diagram showing the differentially transcribed regions: upregulated regions (up, light blue) and downregulated regions (down, violet) over the SOX2 depletion time course. **(G)** Volcano plot showing the downregulated (violet) and upregulated (light blue) transcript after 1 hour of SOX2 degradation. **(H)** Genomic tracks showing ATACseq data (top) and nascent transcription measured with TT_chem_seq (bottom) at the *Platr11* locus in untreated 0.5 and 1h dTAG-13 treated SOX2-FKBP cells. **(I)** As in (H), but showing the accessibility and transcription of the *Hmgcs1* gene.

The increased temporal resolution enabled us to calculate parameters for the dynamics of each individual DAR. To this end we fitted a log-logistic curve describing the decrease or increase in accessibility. From these curves we can derive for every site the moment following depletion at which there is a half-maximal change in accessibility, a value we refer to as T_50_ (Figure 3C). The distributions of the T_50_ values for increases and decreases in accessibility (Figure 3D) confirms the analysis of significantly changed peaks (Figure 3B). The T_50_ values enabled us to rank the peaks that lose accessibility according to their rate of decay. When we determined the motif enrichment as a function of the T_50_ rank we found that the most rapidly changing peaks are enriched for motifs containing the SOX2 consensus motif (Figure 3E). Peaks that rank later according to their T_50_ values show enrichment for KLF2/4 and ZIC2, which could mean that these accessible sites are not directly regulated by SOX2. It is worth noting that our models suggest that the earliest of these putative secondary effects occur already 30 minutes after the induction of SOX2 depletion.

To understand what determines a differentially accessible region (DAR) from a non-DAR (nDAR) we analyzed the protein composition at these sites using a compendium of publicly available ChIPseq data in mESCs (i.e. Cistrome)(Zheng et al., 2019). We reasoned that important factors would be predictive of these sites, so we aimed to discriminate DARs from a matched control set of nDARs using random forest classification. After filtering, we considered 233 unique DNA binding factors as predictors. As a performance metric, these factors reached an area under (AU) the receiver operator characteristic (ROC) of 0.974 (Figure S3A). Unsurprisingly, SOX2 binding was the factor with the greatest importance in the prediction of the DARs. Including only the top 4 chromatin binding factors for prediction was sufficient to perform above an AUROC of 0.90 (Figure S3B). The second most important factor was the BAF (a.k.a. mSWI/SNF) complex member SS18 (King and Klose, 2017), which showed specific binding at DARs (Figure. S3C), hinting at a role for this complex in maintaining open chromatin at sites controlled by SOX2. Next, we wanted to gain insight into the specific histone modifications that are predictive for DARs, so we performed random forest analysis on 38 histone modification ChIPseq datasets (Figure S3A), which reached an AUROC of 0.869. Interestingly, the two histone modifications that are most important in the prediction of differential accessibility are the non-canonical histone modifications H2BK20ac (Kumar et al., 2016) and H3K64ac (Di Cerbo et al., 2014).The latter has been shown to decrease nucleosome stability providing an explanation why these regions are more likely to form accessible regions (Di Cerbo et al., 2014). We then aligned the ChIPseq data from top predictive datasets over the DARs and nDARs after SOX2 depletion (Figure S3C), showing that those factors are enriched at sites that lose accessibility.

Although a pivotal role of SOX2 is to facilitate the establishment and maintenance of nucleosome free regions, not all SOX2 bound regions are necessarily OCRs (Figure S4A,B, Bound CCR: Bound Closed Chromatin Regions). Furthermore, although our results show that SOX2 binding is important for DARs (Figure S3A), SOX2 occupies over 9,000 OCRs that show no change in accessibility following SOX2 depletion (Figure S4A,B bound OCR). Finally, in spite of the fact that the majority of DARs at 2hpd are bound by SOX2, there is a sizeable fraction of sites (n=1,958) for which no detectable binding could be determined by ChIPseq (Figure S4A, unbound DAR). To characterize these unoccupied DARs we performed motif enrichment and found that the compound OCT4::SOX2 motif and SOX motifs was found in a higher fraction of peaks compared to the unbound OCRs (Figure S4C), suggesting that these unoccupied DARs are nevertheless a consequence of SOX2 activity. An explanation of the lack of significantly detected occupancy could be that the dwell time of the protein at specific sites is too short to be detected by ChIPseq.

### Changes in chromatin accessibility are directly related to transcriptional changes

As a transcription factor, SOX2 is responsible for maintaining transcription of genes associated with the pluripotent state. Because RNAseq measures the stable transcript pool, rapid changes in transcription may be occluded in the total level of mRNA. Therefore, to determine direct effects on transcription we measured the production of nascent transcripts using TT_chem_seq (Gregersen et al., 2020). In TT_chem_seq, pulse-labeling with a nucleotide analog enables the purification of nascent transcripts, allowing the detection of transcription from both genic and intergenic sequences. Already after 30 minutes of SOX2 depletion we find that hundreds of transcripts are significantly de-regulated (Figure. 3F). An example of a gene that is immediately lost following SOX2 depletion is the non-coding pluripotency transcript *Platr11* (Figure 3G,H). Upon closer examination we found that accessibility surrounding the putative TSS of this transcript is also severely diminished following SOX2 depletion (Figure 3H). These results are consistent with SOX2 controlling chromatin accessibility to promote the expression of its target genes. It should be noted that, in contrast to DARs, a roughly equal number of genes show a decrease and increase in transcription upon SOX2 depletion (Figure 3G,H). An example of a gene that increases transcription is *Hmgcs1*, which is initially upregulated, but reverts to pre-depletion levels at 6 hpd (Figure 3I). The regulatory landscape surrounding this gene shows no significant changes in accessibility (Figure 3I).

To systematically determine the relationship between changes in chromatin accessibility and gene expression we aligned the positions of DARs to the TSSs of downregulated genes at 2hpd compared to a set of (expression matched) control genes that do not change expression. We found a strong enrichment of DARs around the TSS of downregulated genes which was not observed for OCRs that do not change in accessibility (Figure 4A). In order to identify the most critical factors that determine which genes are activated by SOX2, we aimed to predict which genes go down in expression upon SOX2 depletion in comparison to an expression matched control set. In absence of a precise enhancer-target map, we exploited the pattern of DAR enrichment around TSSs of downregulated genes for making predictions. Specifically, we used logistic regression using differential accessibility after SOX2 depletion with the gene outcome (downregulated or undetectable change) as dependent variable and the counts (or sum of weights) of different categories of genomic loci as predictors. To ensure our prediction model is not dependent on a single set of control genes we generated 100 samples of expression matched genes. We tested a variety of association rules for predictive performance and found that setting a simple distance threshold from the transcription start site outperforms more complex association rules (Figure S5A). Reasoning that counting peaks using a distance threshold is equivalent to weighting the peaks according to a uniform kernel centered at the TSS, we next explored whether other kernel functions could improve prediction (Figure S5B). The heavy-tailed Laplace or Cauchy probability density functions (PDF) yielded the best predictions, which reflects the notion that enhancers can regulate genes over a long range of linear genome. We continued using the (truncated) Cauchy PDF with the half-maximum weight set to 20 kb for further assessment of predictions (Figure 4B), as the cross-validation error was minimal at this distance. In the end we achieved an AUC of 0.83 (Figure. 4C, S5B) in predicting downregulated genes showing a tight connection between chromatin accessibility and gene activation.

**Figure 4.**
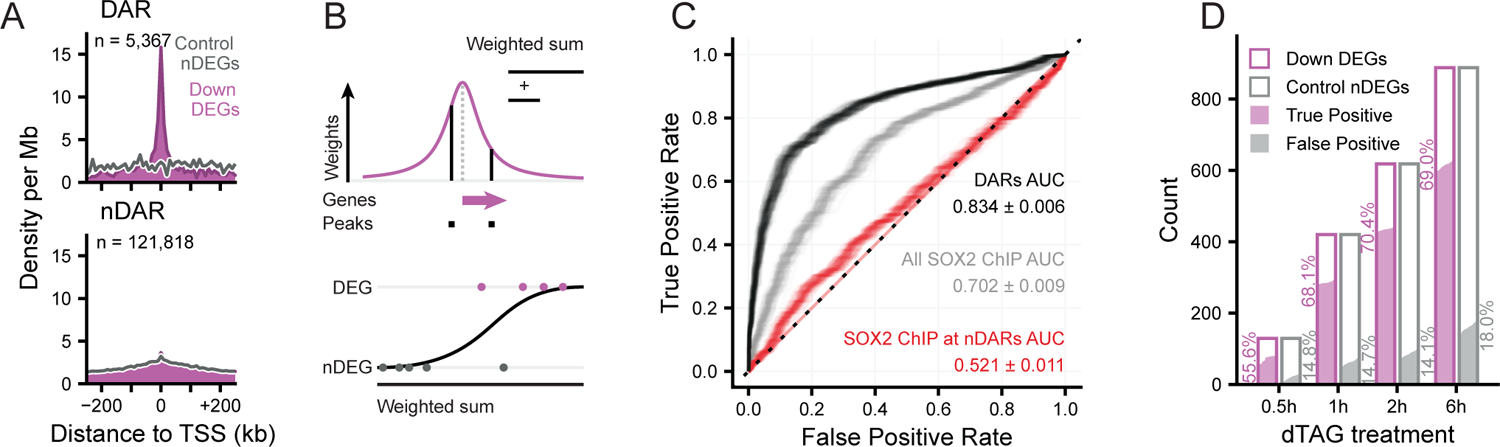
Open chromatin regions maintained by SOX2 are associated with controlling transcription. **(A)** Density of peaks in 10kb bins nearby transcription start sites (TSS) of downregulated (purple) and expression-matched stable control (grey) genes. Panels indicate the set of peaks that were aligned to the TSS: DAR, non-DAR.**(B)** Schematic representation of taking the weighted sum of peaks per gene, weighted by a kernel function (top) and logistic regression predicting the differential expression status of a gene (bottom). (n)DEG = (non) differentially expressed gene. **(C)** Receiver operator characteristic (ROC) curves indicating the predictive strength on differential expression status of downregulated DEGs versus matched control nDEG, for various sets of predictors. The ‘DARs’ set of predictors use ATACseq data. The ‘All SOX2 ChIP’ set has weighted sums of SOX2 ChIPseq peaks. The ‘SOX2 ChIP at nDAR’ has weighted sums of the SOX2 peaks with DARs excluded. Different transparent lines indicate 100 re-samplings of the expression-matched nDEGs. **(D)** Barplot showing the number of downregulated DEG (case) and nDEG (control) genes identified as false or true positive in the predictions with the ‘DARs’ features at the indicated time points. Filled area indicates complementary cumulative distribution function of counts under 100 re-samplings of the expression-matched nDEGs. Percentages indicate the average predictions of the different samplings.

Since SOX2 is crucial is creating OCRs, we wanted to test how differential accessibility compares to SOX2 occupancy in predicting gene expression changes. To test this, we used publicly available ChIPseq data for SOX2 (Marson et al., 2008) in our analyses (Figure 4C). Importantly, solely using SOX2 occupancy as predictor performs starkly worse, showing that accessibility changes are better predictors for genes regulated by SOX2 than occupancy of SOX2 (Figure 4C, S5C). Note that this is not particular to this specific dataset, rather, this is the case for many of the publicly available SOX2 ChIPseq datasets (Figure S5D). Next, we analyzed how well genomic regions bound by SOX2, but which show no change in accessibility following SOX2 depletion, i.e. where SOX2 displays no pioneer activity, could predict expression changes (Figure 4C). Remarkably, these regions had essentially no predictive power for changes in gene expression (Figure 4C, S5C). From these results we conclude that the genomic regions where SOX2 acts as a dominant pioneer factor are crucial for SOX2 mediated gene activation. The sites that are bound by SOX2, but where no it exerts no pioneer activity are largely inert with respect gene regulation.

Next, we decided to determine the predictive power of accessibility changes at different timepoints. Data derived from 2hpd samples were most predictive, wherein we can correctly predict downregulation of about 70% of genes on average, at the expense of incorrectly flagging about 14% of non-DEGs as downregulated (Figure 4D). We suspect that prior to 2 hours the prediction is hindered by the time it takes to establish and detect the primary effects, whereas after 2 hours secondary effects may start to weaken the causal linkage between accessibility and transcription. These results show that TF depletion followed by nascent transcription mapping is a powerful method to prioritize direct from indirect regulatory targets of a TF. Furthermore, our results show that chromatin accessibility is tightly linked to transcription and, in fact, differential accessibility has more predictive power than ChIPseq data with regard to which sites drive gene expression.

### Disruption of a SOX2 dependent OCR confirms prediction of a putative regulatory element

We have demonstrated that SOX2 depletion alters the expression of many genes and that changes in chromatin accessibility can predict which genes respond to SOX2 depletion. Although this strongly suggests that these accessible regions directly control the expression of their putative target genes, the results remain correlative. Based on our prediction model the gene that has the highest probability of being affected by SOX2 depletion is *Klf2*. Correspondingly, we find that *Klf2* transcription is strongly reduced, already after 30 minutes (Figure 5A). Flanking the TSS of *Klf2* there are two OCRs that rapidly lose accessibility after SOX2 depletion. One is 3kb upstream from the TSS and the second is 15kb downstream of the TSS; both are bound by SOX2. Distal regulatory elements are thought to regulate genes through physical proximity within the space of the nucleus (De Laat and Duboule, 2013). To determine whether the regulatory elements around the *Klf2* gene are interacting with the *Klf2* promoter we made use of Micro-Capture-C (MCC), a recently developed basepair resolution chromosome conformation method (Hua et al., 2021). Our analyses show that both the proximal and the distal enhancer interact with the *Klf2* promoter, further suggesting that these elements are functional in driving *Klf2* gene expression.

**Figure 5.**
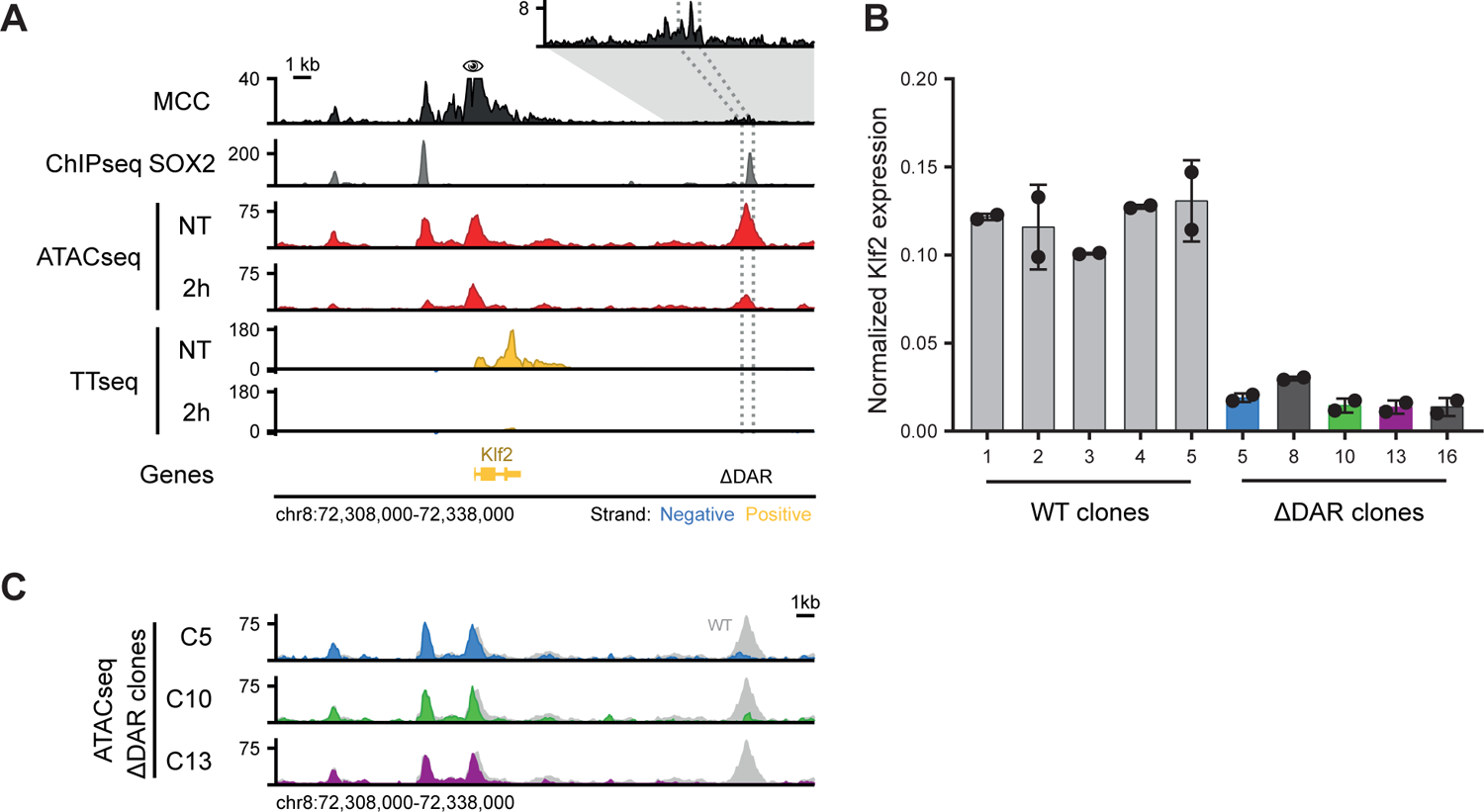
*Klf2* transcription is dependent on a downstream regulatory element that is in physical proximity with the promoter. **(A)** Genomic tracks showing physical interactions with the *Klf2* promoter measured by micro-capture-C (MCC) and SOX2 binding measured by ChIPseq. ATACseq and by TT_chem_seq data is shown for untreated and 2 hours dTAG-13 treatment in SOX2-FKBP cells. The eye symbol indicates the viewpoint for MCC. Contacts with a downstream DAR is shown in the zoomed-in inset. Dotted lines indicate the region targeted for deletion. Y-axes indicate reads per genomic content (RPGC). **(B)** Barplot showing *Klf2* expression in clones with intact (WT) and disrupted DAR (ΔDAR) indicated in (A), as measured by RT-qPCR normalised to the *Rsp26* housekeeping gene. Bar heights indicate means and error bars indicate standard deviation for 2 replicates. **(C)** Genomic tracks showing ATACseq signal around the *Klf2* locus upon knockout of the DAR indicated in (A) in three ΔDAR clones. In grey, accessibility of the NT sample in (A) is shown for comparison. Y-axes indicate RPGC.

In order to demonstrate that our prediction of functional regulatory elements based on differential ATACseq and TT_chem_seq is correct, we wanted to genetically address the causality. We used CRISPR-Cas9 genome editing to homozygously disrupt the downstream putative regulatory element (Figure S6A,B) and selected 5 clones in which the putative regulatory element was disrupted. In these clones, and a corresponding set of wild-type clones, we measured *Klf2* expression levels using RT-qPCR (Figure 5B, S6C). In all the mutant clones the level of *Klf2* was drastically decreased, by ∼85% on average, showing that the region indeed serves as a cis-regulatory element that enhances gene expression. Because the contribution to expression of this one DAR was rather large given multiple nearby DARs that purportedly also contribute, we wondered whether the downstream DAR might act synergistically with the upstream DARs. If this were the case, we reasoned that we might see a concomitant loss in accessibility at the upstream DARs in the lines where the downstream DAR is disrupted. To test this, we performed ATACseq in three of the mutant clones (Figure 5C). We could detect hardly any change in accessibility of the upstream elements in the clones in which the downstream elements were disrupted. Therefore, we conclude that accessibility of the upstream regulatory region is autonomous from the downstream element.

Because ATACseq on the edited clones showed some residual ATAC signal at the DAR, this suggested that the genomic region was not fully deleted. To determine the nature of the disruption of the DAR we PCR amplified a 2.3kb region overlapping the DAR and performed paired-end Illumina sequencing (Figure S6D). This enabled us to assemble to locus following genome editing. For clone 13 we observe a deletion, which is consistent with the lack of sequence reads in the ATACseq. In clone 5 we observed a slightly smaller deletion, but the ATACseq still detects sequence reads in this locus which suggests that the allele is still found in the genome, but was not amplified in our PCR. Interestingly, clone 10 shows a complex pattern of rearrangements following the genome editing. One allele in this clone still harbors an intact SOX2 binding motif region intact, while there is some rearrangement to the upstream sequence. This suggest that, not only the presence of SOX2 binding motifs, but the integrity of the entire locus is essential for proper enhancer activity. Our genome editing results show that using inducible depletion of a pioneer factor coupled to nascent transcription and chromatin accessibility mapping enables us to determine functional cis-regulatory elements that are necessary for transcription.

### SOX2 pioneer activity depends on the DNA binding domain and the transactivation domain

In addition to a specialized DNA binding domain (DBD), most transcription factors also contain unstructured domains that give further specificity to the TF. SOX2 binds DNA with its N-terminal high-mobility group (HMG) domain. Interaction of SOX2 with other proteins is promoted mostly by its trans-activation domain (TAD) at the C-terminus (Cox et al., 2010). Our degron lines offer an opportunity to investigate the role of the different protein domains in the opening of chromatin in the native chromatin environment in living cells. Degron lines have previously been employed to perform separation of function assays (Nora et al., 2020). By ectopically expressing truncated proteins followed by acute depletion of the endogenous protein one can assess the effect on a trait of interest, such as the establishment of chromatin accessibility. We designed expression plasmids that expressed full-length or truncated versions of the SOX2 protein fused to GFP, or GFP alone as a control (Figure 6A). We transfected the plasmids encoding the ectopic fusion proteins (Figure 6B) and monitored their expression level, correct size and nuclear localization (Figure 6B,C) to ensure their functionality. We then treated the transfected cells with dTAG to deplete the endogenous SOX2 protein or DMSO as a control and performed ATACseq on the sorted cells (Figure 6D). As expected, expression of the GFP protein alone resulted in a loss of accessible sites similar to the untransfected cells (Figure 6E). Expression of the full-length SOX2-GFP fusion did not show dramatic loss of accessible sites, indicating that an ectopically provided SOX2-GFP fusion protein can exhibit pioneering activity similar to endogenous SOX2 (Figure 6E). Cells that express the SOX2-HMG-GFP fusion protein, however, were not able to rescue the loss of accessibility observed following SOX2 depletion. Similarly, cells that express the SOX2-TAD-GFP fusion protein were also unable to rescue the loss of accessibility. Our results show that the SOX2 DBD is not enough to promote the formation of accessible sites, rather it is the concerted action of both the DBD and the TAD that is required to establish an open chromatin environment.

**Figure 6.**
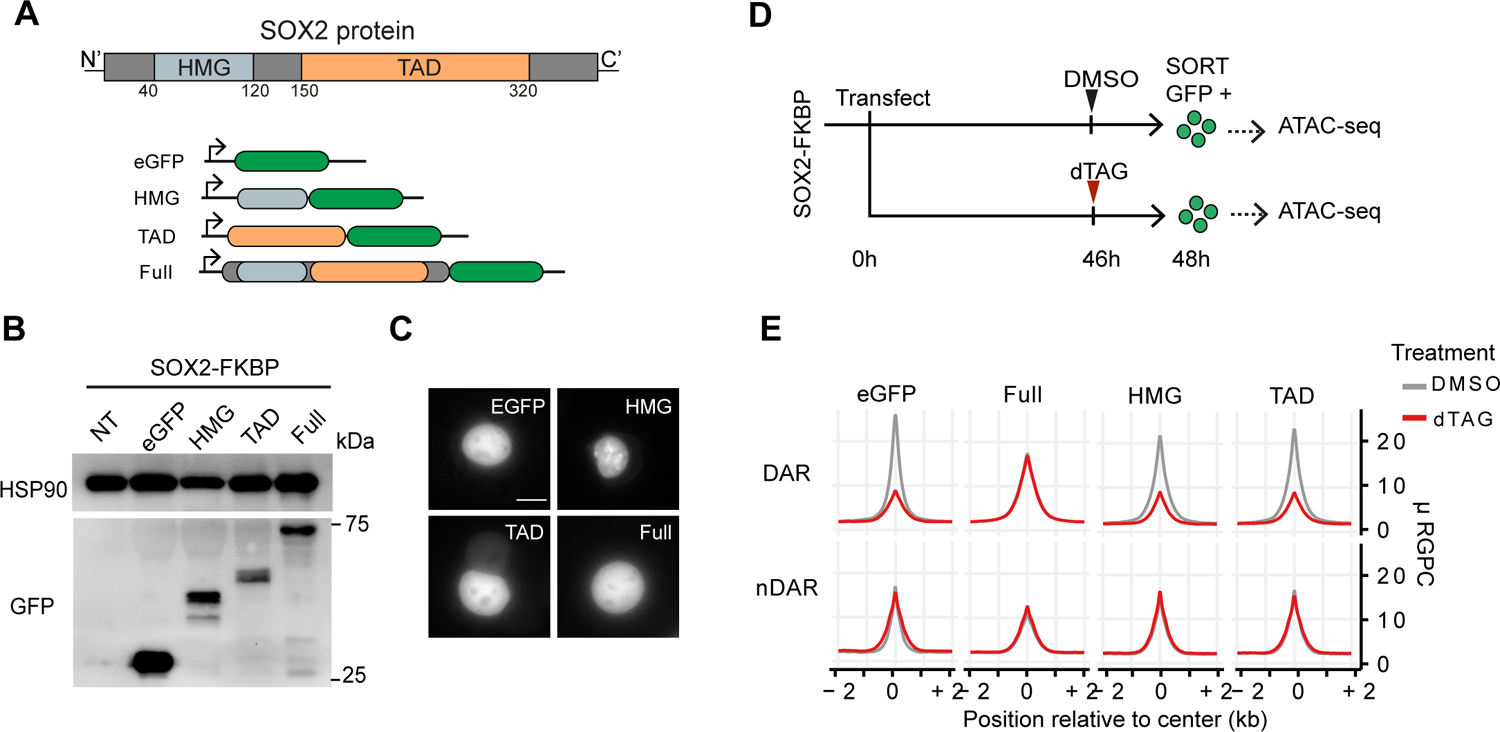
Both the DNA binding and trans-activation domain of SOX2 are needed for pioneer activity. **(A)** Schematic representation of SOX2 protein showing the DNA binding domain (HMG) and the trans-activation domain (TAD). To generate versions of the protein for ectopic expression, the HMG, TAD or full length SOX2 was cloned in frame with the EGFP sequence in an episomal expression vector. **(B)** Western blot using an antibody against EGFP shows the fusion proteins running at the correct size. HSP90 was used as a loading control. Scale bar 10 μm. (NT; Not transfected) **(C)** Representative image of the live EGFP signal and its nuclear localization in transfected cells. **(D)** Representation of the experimental procedure for ATACseq after ectopic expression of the truncated SOX2-EGFP constructs. SOX2-FKBP cells were transfected with the plasmids and seeded for DMSO or dTAG-13 treatment. GFP positive cells were sorted and ATACseq was performed on the EGFP expressing cells. **(E)** Line plot showing the average signal of ATACseq after overexpression of the plasmids after DMSO (grey) or dTAG13 treatment (red) for the endogenous SOX2-FKBP degradation.. The average ATACseq signal is plotted for the DAR and nDAR.

## Discussion

Here we use acute protein depletion to determine the dynamics of chromatin remodeling induced by pioneer transcription factors. We find that loss of SOX2 and OCT4 leads to a rapid loss of accessible sites in the genome. For SOX2 we find that already within half an hour of depletion over a thousand open chromatin sites are significantly decreased in accessibility. This is consistent with a previous study that showed rapid loss of accessibility following OCT4 depletion (Friman et al., 2019). Our observations are also in line with recent reports that used catalytic inhibitors of the SWI/SNF complex or a molecular degrader of SWI/SNF complex member BRG1 that showed that loss chromatin remodeling activity results in a rapid change in the open chromatin landscape(Iurlaro et al., 2021; Schick et al., 2021; Xiao et al., 2022). This indicates that continuous activity of both the transcription factors and these complexes is required for the maintenance of open chromatin. Our results show that maintenance of open chromatin is a highly dynamic process. The fast and direct effects on chromatin and gene expression emphasize the importance of studying the early molecular events that are in play that can lead to transition to a different cell state.

### Open chromatin sites are highly predictive for gene expression dynamics

One of the challenges of assigning functionality to the regulatory landscape of cells is that the number of putative regulatory sites is often much larger than the number of genes. How these regulatory elements collaborate to drive gene expression is an important unresolved question. We have developed a statistical framework to predict changes in transcription based on changes in chromatin accessibility. We show that integrating acute protein depletion methods with direct read-outs of chromatin features (e.g. ATACseq) and transcription (with TT_chem_seq) is a powerful combination to identify functional accessible chromatin sites. The highly dynamic nature of the open chromatin sites means that non-acute methods will be much less accurate in predicting the transcriptional changes, because indirect effects start to play an important role once secondary regulators are up- or downregulated. An example of this is the *Klf2* gene, which encodes for a transcription factor and rapidly loses expression following SOX2 depletion and may in turn regulate other genes.

We find that the highest predictive power is attained when we use a function where weights are assigned in a decreasing manner from the TSS of a gene. This seems to mimic the decreasing contact frequency that is observed as a consequence of polymer dynamics of the chromatin fiber (Zuin et al., 2022). Although the optimal predictive power is reached when we use a distance of 20kb, we would like to note that our analyses do not rule out regulation over larger distances. However, our results strongly suggest that the majority of gene regulation occurs relatively close to the promoter of a gene. Activation of genes by enhancers over larger distances may depend on the presence of additional regulators such as CTCF and cohesin (Kubo et al., 2021; Rinzema et al., 2021). A complicating factor in predicting regulation over larger distances is that although it may result in the correct prediction of genes that are dependent on a particular long-distance enhancer, it will also result in an even stronger increase in false positive predictions. How the cell can make the correct decision to upregulate the correct gene is an important question for the future. One possibility is for instance promoter enhancer compatibility (Martinez-Ara et al., 2022), which is something that we cannot easily explore in our system.

Our results clearly show that pioneer activity of a transcription factor is a stronger predictor for changes in transcription than transcription factor binding on its own. In fact, when we use SOX2 binding sites where we detect no changes in accessibility (either because they are not accessible or do not change in accessibility) to predict gene expression changes we find that these have effectively no predictive power. This has important implications for how to interpret genome-wide binding data, particularly for pioneer factors. SOX2 pioneer activity is crucial in maintaining gene expression, but SOX2 is largely dispensable for regulating transcription at sites where it does not perform its pioneer function. How pioneer factor activity and the formation of accessible sites contribute to gene expression is not fully understood. It may create binding sites for secondary transcription factors (Isbel et al., 2022) which, in turn, recruit co-factors such as Mediator (Allen and Taatjes, 2015). The compendium of genomic regions that are directly controlled by pioneer factors that we describe here can serve as a starting point for further investigating these questions. The degron lines that we have developed are a powerful tool for these future research questions. What the function of SOX2 binding at non-pioneer sites is —if any— is an interesting question for the future, although our results suggest that SOX2 binding alone is not directly involved in gene expression regulation.

Following the genetic disruption of the upstream *Klf2* regulatory element only 15% of Klf2 expression remains. We identified two elements flanking the *Klf2* gene that both lose accessibility following SOX2 depletion. We can consider two models for the regulation by these elements, an additive and a super-additive model. In the additive model gene expression is the sum of activity of the flanking regulatory elements. This would mean that the distal downstream element drives 85% of expression and the proximal element a much smaller part of expression. Massively parallel reporter assays show that the both elements drive expression in roughly equal measure in an episomal context (Martinez-Ara et al., 2022). An alternative is the super-additive model in which multiple weak enhancers synergistically activate gene expression to a higher level than would be predicted from the sum of all enhancer activities. This behavior has been observed for in the *Fgf5* locus (Thomas et al., 2021). Further genetic dissection of the *Klf2* locus is required to determine the mode of action in this locus.

### A model for SOX2 dependent regulatory element establishment

We have shown that not all SOX2 binding sites that are accessible lose accessibility and our computational analyses give insight into which additional factors assist SOX2 in maintaining the accessible state. Apart from SOX2, SS18 is the most predictive protein for differential accessibility following SOX2 depletion. Our separation of function assays showed that SOX2 needs both its DNA binding domains and the transactivation domain for the formation of open chromatin. Based on our data we propose the following model: SOX2 binds to its cognate binding site, which can be the compound site with OCT4 or the sites where it binds alone or with a different factor. Following binding, it recruits the SWI/SNF complex through its transactivation domain, resulting in the remodeling of the nucleosomes around the binding site. Furthermore, the genomic regions that are kept in an accessible state by SOX2 are enriched for acetylated histone modifications such as H2BK20ac, H3K64ac and H3K27ac. The latter two are known to be a consequence of the activity of the p300 acetyltransferase (Di Cerbo et al., 2014; Dancy and Cole, 2015), but whether p300 is directly recruited by SOX2, a secondary consequence of SOX2 binding or recruited through a parallel mechanism remains to be determined. Acetylation of histones is thought to change the electrostatic interaction of DNA with the histone octamer which could assist the nucleosome remodelers in creating accessible sites as a direct consequence of SOX2 binding. An elegant recent study examined the remodeling of nucleosomes by Nanog, Pouf5f3 and Sox19b during zygotic genomic activation in zebrafish. These orthologs of the reprogramming factors included in our study were able to remodel nucleosomes more efficiently when nucleosome occupancy was high, suggesting that the proper positioning of nucleosomes can be a contributor to pioneer activity (Miao et al., 2022). Specific nucleosome positions may arrange the binding site of the pioneer factor such that it can be bound more efficiently (Michael and Thomä, 2021).

Our results emphasize the importance of using acute depletion tools to dissect direct and indirect relationships in the gene regulatory network. Furthermore, they enable the distinguishing functional and non-functional binding sites a specific transcription factor. These two features combined should ultimately lead to improved models for how transcription factors induce gene expression changes as a consequence of developmental cues or external stimuli.

## Acknowledgements

We would like to thank Miguel Martinez-Ara for help with the genomic editing of the *Klf2* enhancer. We thank Richard Young for providing the OCT4-FKBP cell line. We thank the NKI Genomics Core facility, FACS facility, Protein facility, Proteomics facility and Research High Performance Computing facility. We thank members of the de Wit lab for critically reading the manuscript. Work in the de Wit lab is supported by the Dutch Research Council (016.161.316, Vidi) and the European Research Council (865459, ‘FuncDis3D’). M.M., T.v.d.B, H.T. and E.d.W. and are part of Oncode Institute which is partly financed by the Dutch Cancer Society.

## Author Contributions

M.M. performed experiments, analyzed the data and wrote the manuscript. T.v.d.B. analyzed the data and wrote the manuscript. H.L. performed and analyzed MCC experiments. J.O.J.D. oversaw MCC experiments. H.T. performed experiments. E.d.W. conceived and supervised the project and wrote the manuscript. All authors edited and contributed to the manuscript.

## Materials and Methods

### Cell lines and gene targeting

Mouse Embryonic Stem Cells (ESCs) cell lines were cultured on 0.1% gelatin-coated plates in serum-free DMEM/F12 (Gibco) and Neurobasal (Gibco) medium (1:1) supplemented with N-2 (Gibco), B-27 (Gibco), BSA (0.05%, Gibco), 10*4U of Leukemia Inhibitory Factor/LIF (Millipore), MEK inhibitor PD0325901 (1 µM, Selleckchem), GSK3-β inhibitor CHIR99021 (3 µM, Cayman Chemical) and 1-Thioglycerol (1.5×10-4 M, SigmaAldrich). The cell lines were passaged every 2 days in daily culture. E14 mESC were used for NANOG and SOX2 (IB10 mESC) targeting. The OCT4-FKBP (v6.5 mESC) were kindly provided by Richard Young(Boija et al., 2018). For depletion treatment, the OCT4-, SOX2-, NANOG-FKBP cell lines were treated with a final concentration of 500 nM dTAG-13 or DMSO (NT, Not treated). For time course experiments, cells were seeded overnight and treated the next day, by inducing protein depletion at different time points and harvesting at the end of the time course. DMSO was added at the same time as the latest dTAG-13 treatment.

For the knock-in of the *FKBP* sequence at the genes of interest, we used previously described plasmids and approach (SOX2 fkbp-donor, Addgene # 175552. SOX2 sgRNA, Addgene #175553. NANOG donor, Addgene # 175554. NANOG sgRNA, Addgene # 175555) Briefly, cells were transfected with the plasmids containing the gRNA seqeunce and the donor plasmid designed to include the FKBP-2xHA-P2A-[GFP/mCherry] in between two homology arms for the gene of interest. After transfection, cells were sorted in 96 well plates and manually picked for genotyping using PCR and western blot. Homozygous clones responding to dTAG-13 were used for experiments.

### Western Blotting

Cells were harvested and lysed in RIPA lysis buffer (150 mM NaCl, 1% NP-40, 0.5% sodium deoxycholate, 0.1% SDS, and 25 mM Tris (pH=7.4). A 10% SDS-PAGE gels were used to separate the Nanog, Oct4 and Sox2 proteins. Protein was transferred to a pre-activated PVDF membrane using Trans-Blot Turbo Transfer System (Bio-Rad). The blots were incubated with the following primary antibodies overnight at 4 °C: SOX2 (1:1000, D9B8N, Cell Signaling), OCT4 (1:1000, D6C8T, Cell Signaling), NANOG (1:1000, D2A3, Cell Signaling), GFP (1:1000, ab6673, abcam), and as loading control HSP90 (1:2000, 13171-1-AP). After incubation, the blots were washed 3 times with TBS-0.1% Tween-20. The blots were then incubated with secondary antibody against rabbit IgG at room temperature for 1 h, following by 3-time TBS-0.1%-Tween-20. The proteins attached with antibodies were hybridized with Clarity Western ECL Substrate reagent (Bio-Rad) and visualized in a ChemiDoc MP Imaging System (Bio-Rad).

### ATACseq

Cells were harvested and washed 1X in cold PBS. 50,000 cells were counted and nuclei were lysed using 2x lysis buffer (1M Tris-HCl pH7.5, 5M NaCl, 1M MgCl2, 10%IGEPAL). Cells were spun down incubated for 1h at 37°C in tagmentation buffer (20 mM Tris(hydroxymethyl)aminomethane; 10 mM MgCl2, 20% dimethylformamide, brought at pH 7.6 with acetic acid) containing custom Tn5 produced by the NKI in-house Protein Facility. For amplification of the tagmented DNA, two rounds of PCR were used using KAPA kit amplification protocol and sequencing adapters were included during the PCRs for library preparation. Fragments smaller than 700 bp were purified using SPRI beads selection. Quality of the libraries was checked on a Bioanalyzer before sequencing. All ATACseq experiments for NANOG and OCT4 depletion were done in duplicate and for SOX2 we performed experiments in duplicate as well as in an independent biological clone.

### Genome editing of Klf2 regulatory element

For deletion of the *Klf2* downstream regulatory region, SOX2-FKBP cells were used. CRISPR-Cas9 was used to induce the deletion using sgRNAs targeting sites flanking the region. sgRNAs were cloned into a plasmid expressing GFP (Plasmid #111596, Addgene). The unmodified PX330 plasmid (Plasmid #42230, Addgene) was delivered into the cells to express the Cas9 protein. Plasmids containing the sgRNA and Cas9 were co-tranfected into the cells with nucleofection Kit Amaxa (VPH-1001)) following the manufacturer’s protocol using the A030 program. After 2 days, single GFP positive cells were sorted in a 96 well plate precoated with 0.1% gelatin and expanded. Clones were manually picked and expanded for further experiments. Clones were genoptyped for the homozygous disruption using PCR and Sanger was used to validate the disruption with primers spanning the sites of editing.

Clones for the homozygous deletion were expanded and clones from the parental cell were obtained to control for the endogenous variation of *Klf2* gene expression. RNA was isolated in triplicate from cells using RNeasy Kit (Qiagen cat #74106) and genomic DNA was cleared with DNaseI treatment. Purified RNA was retro-transcribed into cDNA with the iScript cDNA synthesis kit (Biorad). Expression of the *Klf2* gene was quantified with RT-qPCR, with primers for *Rps26* and *Klf2* genes using SensiFast No-Rox kit (Bioline). The expression of *Klf2* was calculated using the 2^-ΔΔCt^, normalizing over the housekeeping *Rsp26* expression.

### Amplicon sequencing

To characterize the deletion at the downstream *Klf2* DAR we performed amplicon sequencing. The genomic region corresponding to the region of interest was amplified using PCR using primers amplifying a 2kb region, spanning the putative deletion site. The PCR product was purified using Qiagen PCR purification Kit (cat n. 28106) and the product was tagmented using 0.02 mg/ml Tn5 for 10 minutes at 55 °C. The reaction was stopped by adding 0.02% SDS for 5 minutes at 55 °C. Similar to the ATACseq protocol, tagmented DNA was used as input for library preparation and amplified by KAPA HiFi DNA polymerase, following the manufacturer’s protocol. Libraries size was checked on Bioanalyzer and samples were sequenced on a MiSeq Nano, paired end 250 bp reads length.

### Ectopic expression of SOX2 domain mutants for ATACseq

SOX2 protein sequence (uniprot #P48432) was used as a reference for identifying the HMG (DNA binding domain), TAD (trans-activation domain) and full length of SOX2 protein, and the corresponding mRNA sequence was used to create the truncation of the proteins. The SOX2 endogenous start codon was added to promote expression of the domains. An eGFP cassette was included in the design to be in-frame with the different domains and used for selection of domains expression. The designed constructs were ordered from Twist Bioscience and cloned into an expression vector with the EIF1A promoter. The eGFP only vector was used as control. Plasmids (5 μg/ 10^6^ cells) were transfected into SOX2-FKBP degron cells with nucleofection following the AMAXA protocol and A030 program. For each plasmid, 5 million cells were transfected and then seeded dividing the total volume of cells into two plates pre-coated with 0.1% gelatin. At 46h after nucleofection cells were treated with DMSO or 500 nM dTAG-13 to deplete the endogenous SOX2-FKBP. At 48h cells were harvested and GFP positive cells were sorted and ATACseq was performed directly after sorting.

### TT_chem_seq

Libraries for TT_chem_seq were prepared following a published protocol(Gregersen et al., 2020). SOX2-FKBP cells were seeded and the day after treated with 500 nM of dTAG-13 (0.5hpd, 1hpd, 2hpd, 6hpd) or DMSO as control. Cells were labeled with 2mM 4SU for 10 minutes. Total RNA was isolated and fragmented. The 4SU-biotin labeled RNA was enriched using streptavidin coated MicroBeads. Libraries were prepared using KAPA RNA HyperPrep kits (Roche) using dual indexing adapters. Libraries were sequenced on a NextSeq 550.

### Quantitative mass spectrometry

For quantitative measure of protein abundance after dTAG treatment, cells from the OCT4, NANOG and SOX2-FKBP and parental cells were expanded and treated with either DMSO or 500 nM dTAG-13, for 30 minutes. Cells were harvested and 30 million cells per condition were centrifuged at 500g x5 minutes, washed with PBS and pellet was snap frozen. All samples were prepared in quadruplicate before analysis.

### Micro-Capture-C

Micro Capture-C was performed as previously described (Hua et al., 2021). Cells were fixed with formaldehyde (2% (wt/vol)) for 10 min at room temperature and permeabilised with digitonin (0.005% (wt/vol)). 2-3 ×10^6^ cells were treated with MNase (NEB M0247) ranging from 10-30 Kunitz U in 800 μl of custom buffer (Tris-HCl pH 7.5 10 mM, CaCl_2_ 1 mM) for 1h at 37 °C. Ethylene glycol-bis(2-aminoethylether)-N,N,N′,N′-tetraacetic acid (EGTA) 5 mM (Sigma E3889) was added to quench the reaction. The reaction was centrifuged (5 min, 300 × g) and the supernatant discarded. The cells were resuspended in 1 ml phosphate buffered saline with 5 mM EGTA of which 200 µl was used to measure the digestion efficiency. The remainder was centrifuged (5 min, 300 × g) and the supernatant discarded. Cells were resuspended in DNA ligase buffer (Thermo Scientific EL0013) supplemented with dNTPs (dATP, dCTP, dGTP and dTTP; 400 µM final concentration of each (Thermo Fischer R0191)); EGTA 5 mM; T4 Polynucleotide kinase PNK 200 U/ml (NEB M0201L); DNA Polymerase I Large (Klenow) Fragment 100 U/ml (NEB M0210L) and T4 DNA ligase 300 U/ml (Thermo Scientific EL0013). The reaction was incubated at 37 °C for 2 h followed by 20 °C for 8h using an Eppendorf Thermomixer at 500 rpm. The ligation reaction was centrifuged and the supernatant was discarded. The chromatin was decrosslinked with proteinase K at 65 °C (>2 h) and the DNA was extracted. Double oligonucleotide capture was performed as previously described (Davies et al., Nature Methods 2016). Data were analyzed with a custom analysis pipeline specifically developed for MCC data analysis.

### Data preparation

Sequencing data from ATACseq and TT_chem_seq were mapped to the mm10 reference genome with the bwa program version 0.7.17-r1188 using the command ‘bwa mem-M’(Li and Durbin, 2009). Alignments were filtered with samtools version 1.10 using ‘samtools view -h -b -q 10’ (Li et al., 2009). Alignments were deduplicated using the ‘MarkDuplicates’ command from Picard tools version 2.12.0, using the argument ‘REMOVE_DUPLICATES=truè. For TT_chem_seq data, alignments were split up by strand as described in (Gregersen et al., 2020). We conducted the rest of the analyses in R version 4.0.5 and Bioconductor version 3.12 (Huber et al., 2015). Bigwig coverage tracks were created using the GenomicRanges, rtacklayer and GenomicAlignments R/Bioconductor packages (Lawrence et al., 2009, 2013), scaling coverage to 1x genome size (reads per genomic content).

### Identification of DARs

To identify DARs, peak calling was performed on merged data from timeseries experiments of all three degron lines, using MACS2 version 2.2.7.1 with the arguments ‘-g mm –nomodel –keep-dup all’ (Liu, 2014). Subsequently, the number of overlapping paired-end fragments were counted per peak. Peaks were subsequently filtered by requiring every peak to have more than 10 reads in more than 2 samples in the SOX2 comparisons, and more than 10 reads in more than 6 samples in comparisons with multiple FKBP-tagged cell lines. Differential accessibility analysis was performed by using DESeq2 with the design argument set to ‘∼ timè in the SOX2 comparisons and ‘∼degron + time + time:degron’ in comparisons with multiple cell lines(Love et al., 2014). The ‘nbinomWaltTest()’ function was used with default arguments to estimate peak-wise p-values and effect sizes assuming the null hypothesis that there was no change over time. A peak was considered differentially accessible when the false discovery rate corrected p-value was below 005. For the DAR^SOX2^ DAR^OCT4^ and DAR^OCT4/SOX2^ categories, we used the thresholds indicated in table 1.

**Table 1:** Criteria for DAR categories. FDR = False discovery rate corrected p-value. LFC = log_2_ fold change

| DAR | Contrast |  |  |
| --- | --- | --- | --- |
|  | SOX2 NT vs 2hpd | OCT4 NT vs 2hpd | Interaction |
| SOX2 | LFC < -0.5 | LFC > -0.5 | FDR < 0.05, LFC < 0 |
| OCT4 | LFC > -0.5 | LFC < -0.5 | FDR < 0.05, LFC > 0 |
| OCT4/SOX2 | FDR < 0.05, LFC < -1 | FDR < 0.05, LFC < -1 | FDR > 0.05 |

### Analysis of DNA binding motifs

Motif position weight matrices (PWMs) were taken from the JASPAR 2020 core vertebrate dataset as provided in the JASPAR2020 R/Bioconductor package(Castro-Mondragon et al., 2022). After matching PWMs to corresponding genes and filtering the PWMs on expression of the DNA binding factor, 408 PWMs were retained for further analysis. The presence or absence of a PWM match in a peak was determined using the motifmatchr R/Bioconductor package (Schep, 2020). Enrichment of motifs was determined by comparing DAR category peaks against all peaks that show no significant change in neither the Sox2, Oct4 or interaction contrasts. To calculate log odds ratios, individual logistic regression per PWM was performed with PWM occurrence in a peak as dependent variable, and (i) peak GC content, (ii) total motif presence per peak and (iii) peak membership to the DAR category as independent variables. The regression coefficient for variable (iii) were taken as log_2_ odds ratios and the significance was determined with a Wald test of this coefficient.

### Quantification of accessibility dynamics

A selection of peaks was made by using the DESeq2’s likelihood ratio test ‘nbinomlrt()’ using ‘∼ 1’ as the reduced model and ‘∼ timè as the full model. Peaks that had an FDR-corrected p-value below 0.05 were kept for further analysis. Library size normalised counts per peak were rescaled to a minimum of 0 and maximum of 1. The ‘drc’ R package (Ritz et al., 2015) was used to fit log-logistic curves to the data, which uses the following formula:

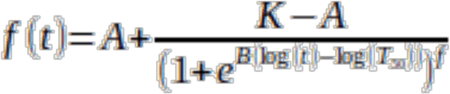

Wherein t is the sampled time, A and K are the lower and upper asymptotes respectively, B is the slope at the inflection point T_50_, and f is also known as the Hill exponent. Multiple parameterizations of models were fit, wherein B and T_50_ were estimated every time, whereas in some models A, K and f were fixed at 0, 1 and 1 respectively. To balance model complexity and goodness of fit between different models, the difference in Akaike’s information criterion (ΔAIC) was used to select the simpler model when the ΔAIC was less than 2. Moreover, we used the ‘lm()’ function to fit the data with three polynomial models of 0th up to 2nd degree. Log-logistic models were preferred over 2^nd^ degree polynomial when the ΔAIC < 2, whereas 0^th^ and 1^st^ degree polynomials were preferred over log-logistic models when ΔAIC < 2. Of the 15,906 DARs that were selected initially, the T_50_ of 9,692 DARs (for which the log-logistic models were preferred) are used in Figure 3.

### Analysis of TT_chem_seq data

To determine transcribed units (TUs), all TT_chem_seq data was pooled per strand, the genome was tiled into 200 bp bins and middle-fragment positions were counted. The tiles were then discretized into low and high transcriptional states with a 2-state hidden Markov model using Poisson-Lognormal distributions with the STAN R/Bioconductor package(Zacher et al., 2020). Runs of consecutive bins of the same state (ignoring single-bin interruptions) were then merged to form putative TUs. Putative TUs were then filtered to have more than 25 overlapping fragments. These TUs were then refined to basepair position by finding positions in a 400bp window where the postive difference between two consecutive coverage positions exceeded a threshold of 1. The putative TUs were then associated to genes in the RefSeq annotation (O’Leary et al., 2016). TUs associated with multiple small genes (such as miRNAs, Jaccard index between TU and annotated gene <0.02) were annotated as such. TUs associated with multiple, larger genes were split when an annotated transcription start site (TSS) internal to the TU was observed to be followed by an increased wave of transcripts. We required these TSSs to be followed by a median increase in coverage of 5 fragments within a 2kb window centered at the annotated TSS, relative to 5’ of the TSS. Subsequently, fragments were counted in the resulting TUs. TUs were further filtered to have more than 10 fragments in at least two samples, after which they were subjected to differential expression analysis with the DESeq2 R/Bioconductor package(Love et al., 2014). ThènbinomWaldTest()’ function with default arguments was used to test the contrasts of treatment time against untreated.

### Prediction of downregulated DEGs

First, various association rules were used to associate peaks with genes. The TSS was taken empirically from the TT_chem_seq data rather than from other annotations. The ‘distance threshold’ rule took every peak with some distance to the TSS of a gene. The ‘basal + extension’ rule associated peaks within a basal domain and in an extended domain (until another gene’s basal domain was encountered)(McLean et al., 2010). The ‘KNN, many TSSs one peak’ rule searched for the k nearest genes for every peak, and the ‘KNN, one TSS many peaks’ searched for the k nearest peaks for every gene. For every gene in consideration, the associated peaks were counted by category (down DAR, nDAR or up DAR). These counts were then used in a logistic lasso regression model to predict a yes/no outcome for gene expression status (DEG or nDEG), using the glmnet R package version 4.1-3(Friedman et al., 2010). Stable control nDEGs were chosen such that their average transcription are similar to the DEGs under consideration and moreover fail to reject the null hypothesis of a Kolmogorov-Smirnoff test that the distributions are equal. Such nDEGs were more numerous than than DEGs, these could be re-sampled to estimate variability in predictions. Appropriate parameters for the association rules were determined by performing a parameter sweep, and choosing a distance, k, extended and basal domains, that minimized cross-validation error. Because a simple distance threshold outperformed other association rules, we explored other distance-based weighting schemes.

Second, for the kernel-based predictions, a 10Mb window around the TSS was used to putatively associate peaks to a gene. Every gene-peak association was then weighted by the distance between the TSS and the summit of a peak according to the kernel functions, which heavily penalise far-distance associations. These were then summarised per gene by taking the sum of weights of all putatively associated peaks for every category of peak. These weighted sums per peak categories were then used as predictors in a similar logistic regression model as described above. Again, appropriate parameters for the kernel functions were found by performing a parameter sweep over various distances and choosing these such that they have minimal cross-validation error. Further predictions were performed for the downregulated DEGs at 2 hours, using the (truncated at 5Mb) Cauchy density function as the kernel function with the ‘scale = 20e3’ argument in R’s ‘dcauchy()’ function, and the down DARs, nDARs and up DARs at the 2h dTAG treatment contrast were used as the predictor categories. Predictions in Figure 4C, S5C and S5D also used Sox2 binding sites as determined by ChIPseq as predictors.

### Random forest classification with Cistrome data

Data for mouse factors and histones were downloaded as batches from the Cistrome Data Browser website (http://cistrome.org/db/). Datasets were enriched from datasets from embryonic stem cells by filtering for the following inclusion and exclusion criteria. Inclusion criteria (i) the cell line was described as ‘V6.5’, ‘E14’ or ‘Mouse embryonic stem cells’, (ii) the cell type was described as ‘Embryonic Stem Cell’ or ‘Stem Cell’ and (iii) the tissue was described as ‘Embryo’. Exlusion criteria were (i) cell types described as ‘Embryonic Cortex’, ‘B lymphocyte’, ‘Embryonic Fibrobtst’, ‘Fibroblast’, ‘Lymphocyte’, ‘Junction cell’, ‘Neuron’, ‘Epithelium’, ‘Hemangioblast’, ‘Mesodermal Progenitor Cell’, ‘Progenitor Cells, Mesenchymal’, ‘Trophoblast Giant Cell’ or ‘Trophoblast Stem Cell’ and (ii) tissues described as ‘Brain’, ‘Colon’, ‘Ectoderm’, ‘Endoderm’, ‘Intestine’ or ‘Liver’. Moreover, datasets were filtered out that had less than 1000 peaks in standard chromosomes, or that had no or multiple associated factors or histones (notably, FAIREseq data and a single H3K9ac, H3K14ac combined ChIPseq dataset). To filter for redundant factors, the Jaccard index was calculated between the dataset and the whole of our peakset (blind to DARs), and per factor, datasets with the highest Jaccard index were kept for further analysis. After this filtering, we separately considered 233 factor datasets and 38 histone datasets as predictors for random forest classifications using the randomForestSRC R package(Ishwaran and Kogalur, 2022). The independent variable to predict was the downregulated DAR status of peaks, among matched control nDAR peaks. Variable importance with confidence intervals were calculated by subsampling inference, and reported variable importance is the normal-Z subsampling confidence intervals with delete-d jack-knife estimator.

### Oligonucleotides

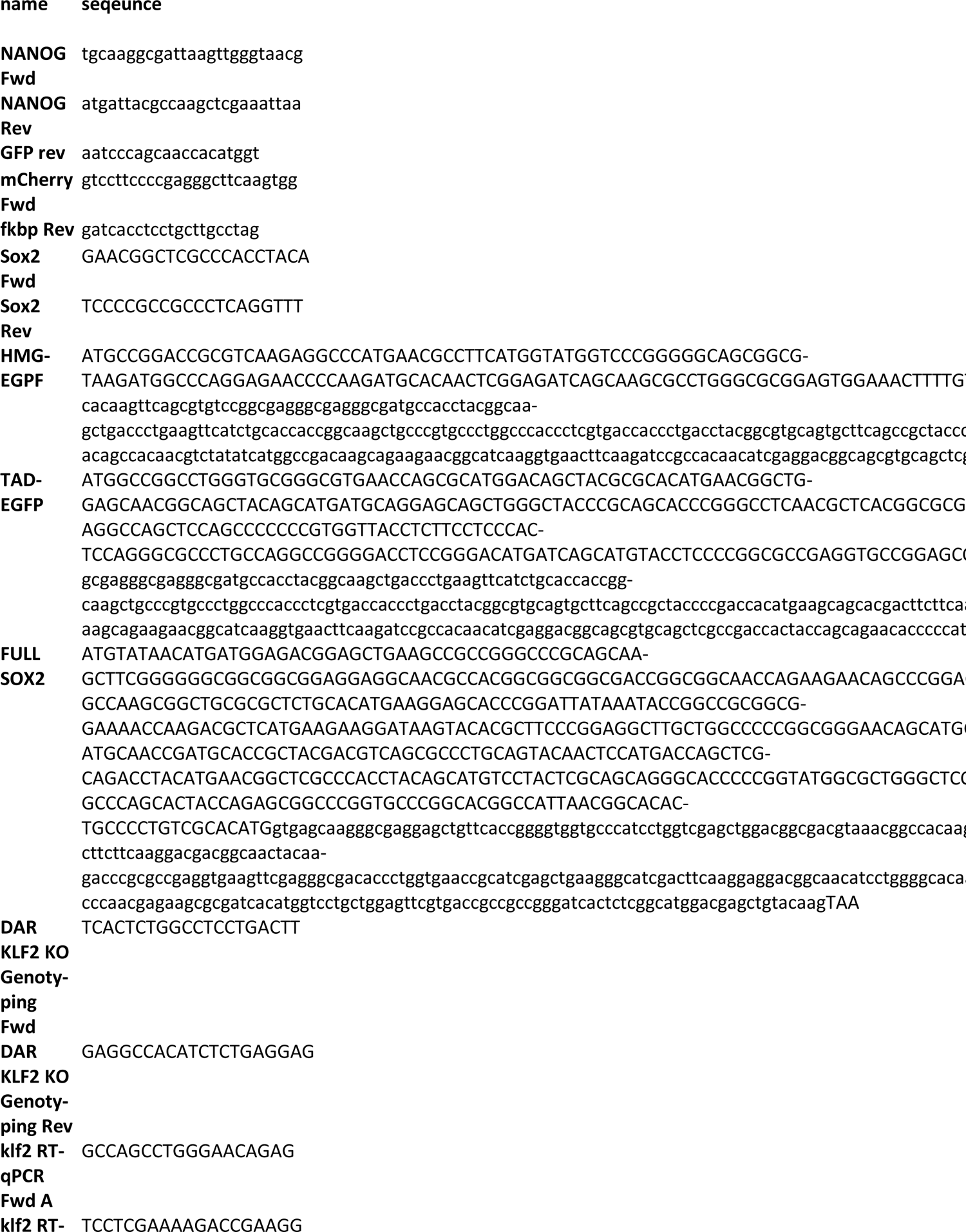
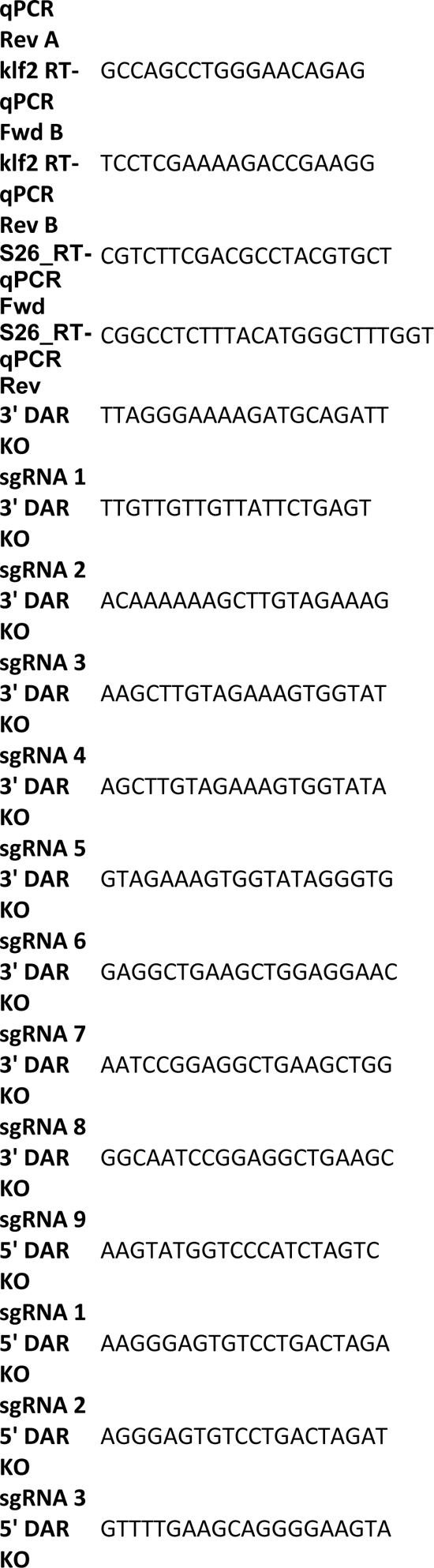
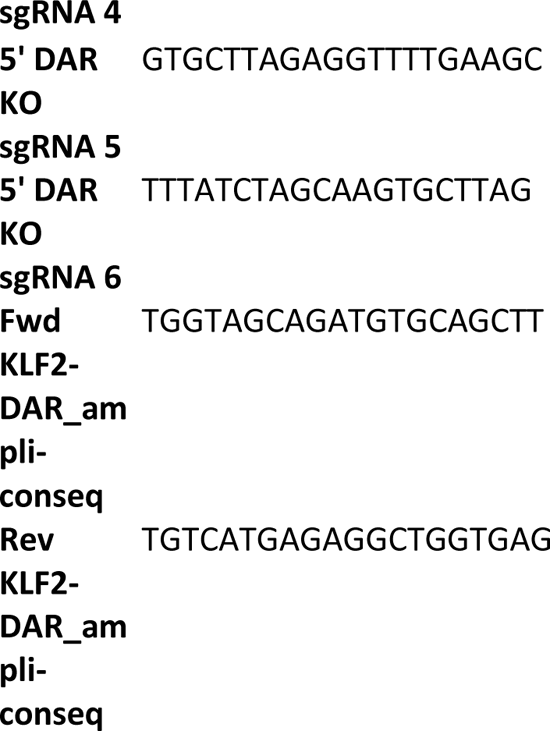

---

**Figure S1.**
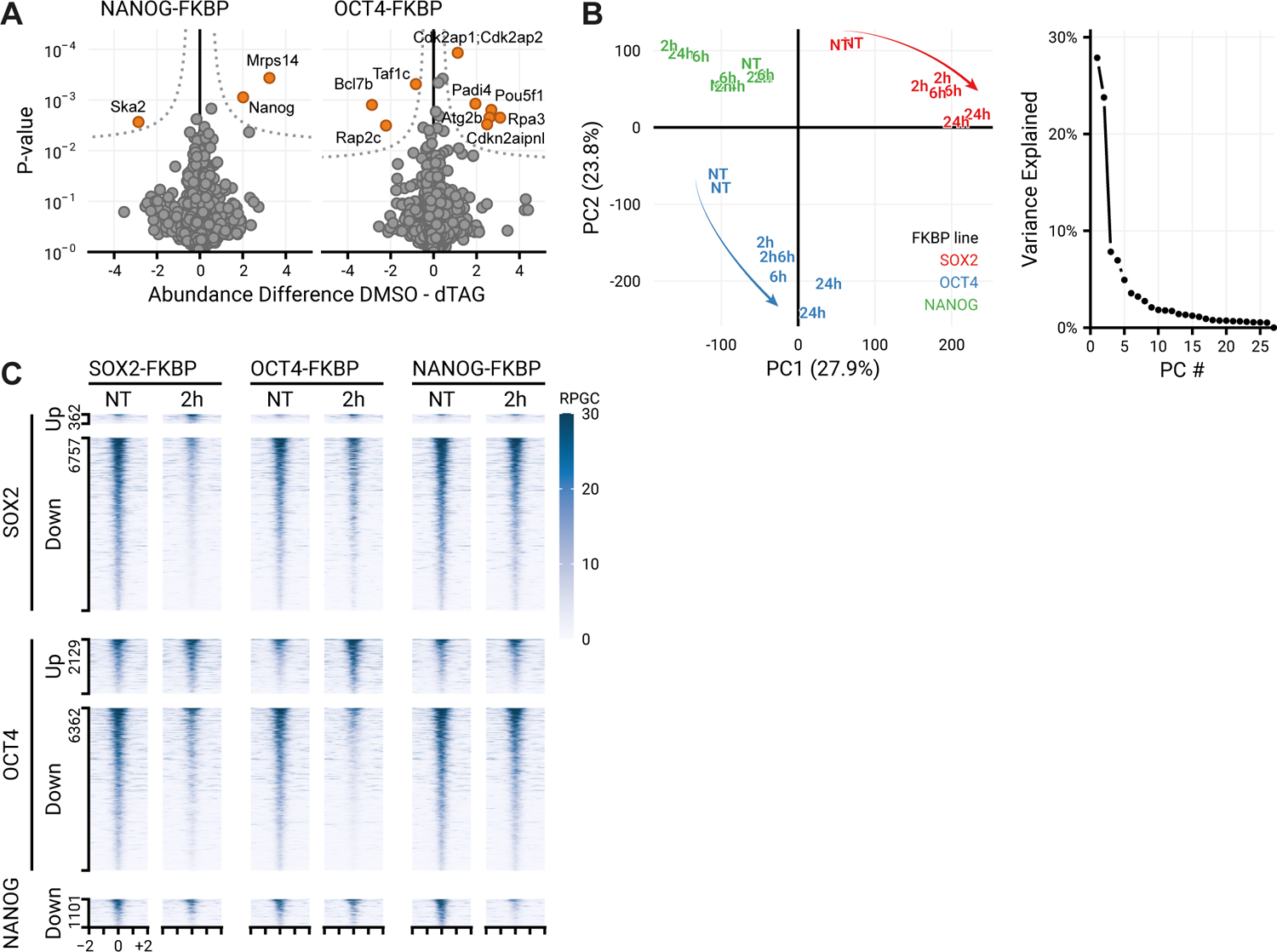
Quantification of changes the proteome and open chromatin landscape following depletion of pluripotency transcription factors. **(A)** Quantitative mass spectrometry results showing the differential protein abundance comparing 30 minutes of dTAG-13 treated and untreated NANOG-FKBP and OCT4-FKBP cells. **(B)** Principal component analysis (PCA) scatter plot of accessibility data of the three FKBP-tagged lines indicating the first two principal components (PCs) (left). Arrows indicate major movements over time. On the right, the amount of variance explained by each principal component. **(C)** Tornado plot showing accessibility in the SOX2-, OCT4- and NANOG-FKBP-lines in untreated cells and following 2 hours of dTAG-13 treatment at differentially accessible regions for the indicated lines. Note that the ‘Up’ category in the NANOG-FKBP has been omitted for having too few peaks.

**Figure S2.**
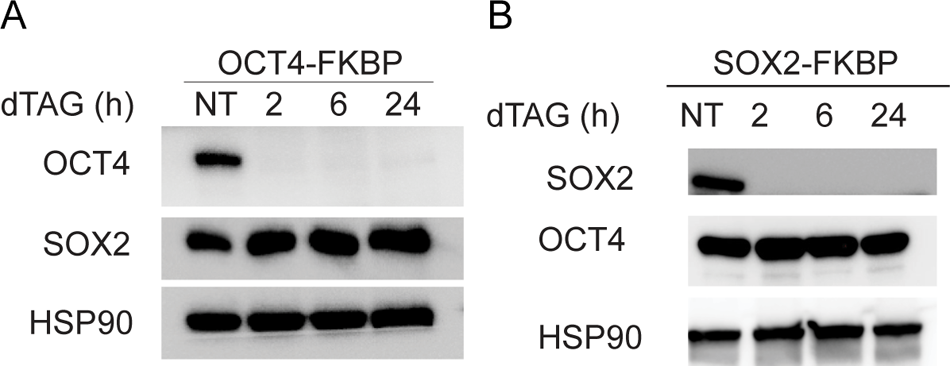
OCT4 and SOX2 depletion does not affect protein levels of SOX2 and OCT4. (A) Western blot showing the levels of OCT4 and SOX2 protein in the OCT4-FKBP cell line after 2, 6 and 24h of dTAG-13 treatment. HSP90 used as loading control. (B) The same as in (A) but in the SOX2-FKBP cell line.

**Figure S3.**
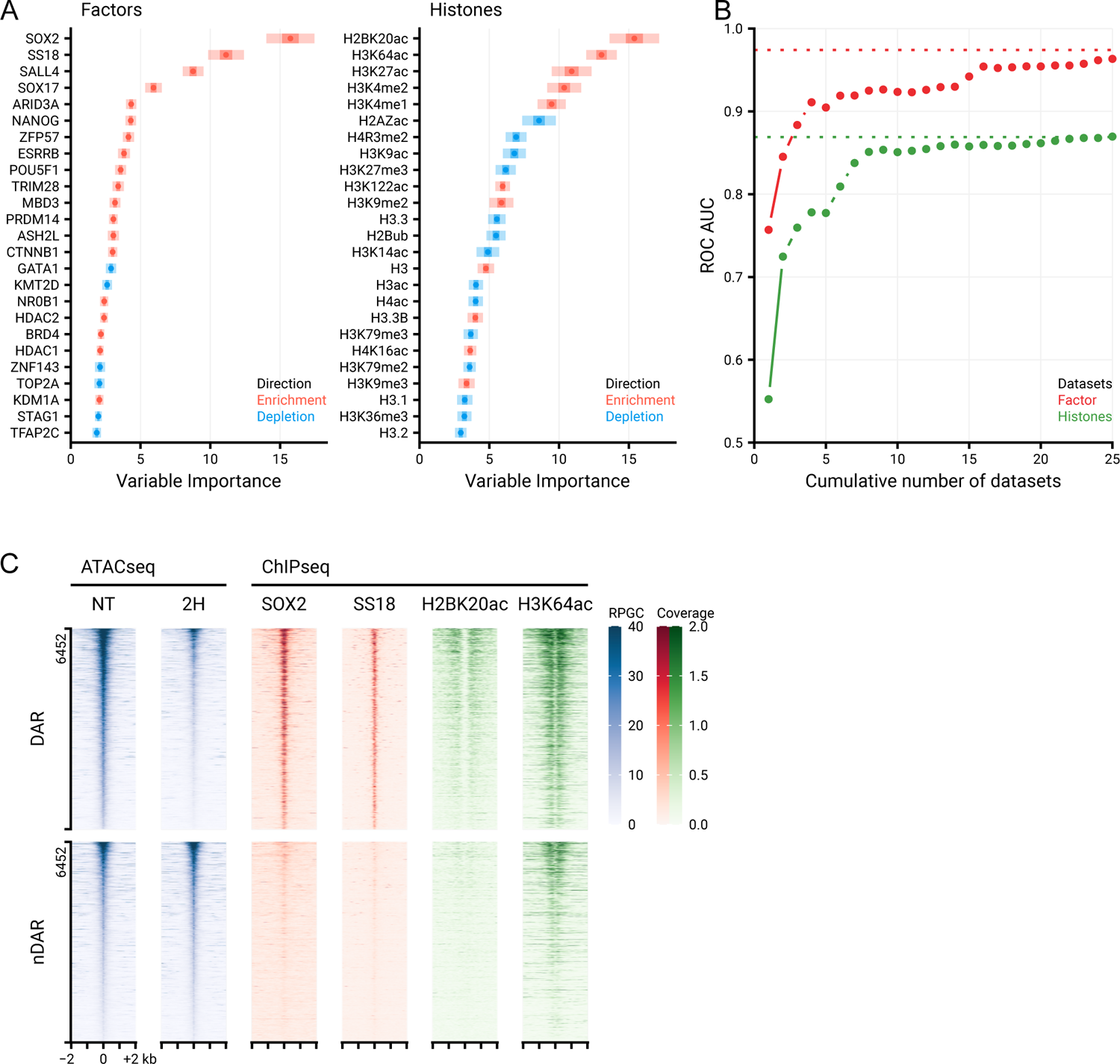
Random forest classification using Cistrome data reveals proteins and histone modifications that can predict differential accessibility following SOX2 depletion. **(A)** Left, top 25 (chromatin binding) factors in the Cistrome factors datasets whose overlap with all ATACseq peaks is predictive in random forest classification to discriminate the DAR from nDAR peaks. ATACseq peaks were extended by 300bp in both directions. Variable importance was calculated with subsampling inference, wherein the 95% confidence interval (CI) is indicated with a light colour, the 50% CI with a darker colour and the median with a point. Enrichment and depletion indicate higher average overlap in the DAR than nDAR categories respectively. Right, top 25 histone modifications using the Cistrome histone datasets.. **(B)** Area under the receiver operator characteristic curve indicating predictive performance using an increasing number of factors/histone modifications sorted by variable importance in discriminating DAR from nDAR peaks. Dotted lines indicate predictive performance of using all factors/histones for prediction (n=233/n=38). **(C)** Tornado plots showing ATACseq data for untreated and 2h dTAG-13 treated SOX2-FKBP cells and ChIPseq signal for the top two predictive datasets from the factors and histone databases over the DAR and nDAR peaks. RPGC = reads per genomic content, coverage indicates values in pre-processed data(Zheng et al., 2019).

**Figure S4.**
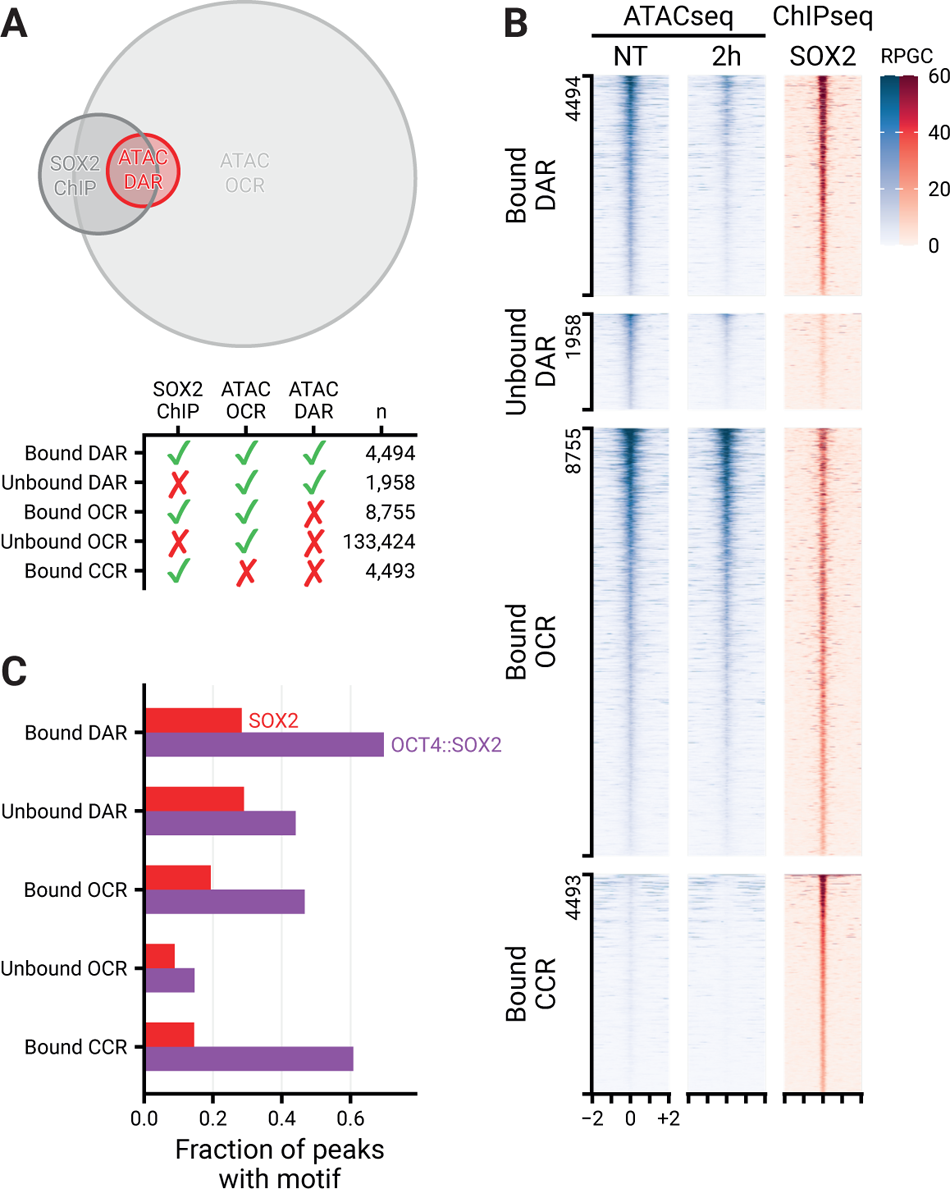
Quantitative and qualitative analysis of ATACseq and ChIPseq. **(A)** Venn diagram showing the overlap between differentially accessible regions (DARs), all ATACseq peaks (OCR) and SOX2 DNA binding (SOX2 ChIP). Bottom panel shows the number of peaks in each overlap category. **(B)** Tornado plot showing accessibility before and after 2 hours of SOX2 depletion, along with SOX2 binding, for the sites in Figure 4A. **(C)** Fraction of peaks containing the Oct4::Sox2 or Sox2 DNA binding motifs, stratified by whether they are accessible peaks as measured by ATACseq, SOX2 binding sites as measured by ChIPseq, differentially accessible after 2 hours of SOX2 depletion or combinations thereof. (OCR, Open chromatin regions. DAR, Differential accessible regions. CCR, closed chromatin regions)

**Figure S5.**
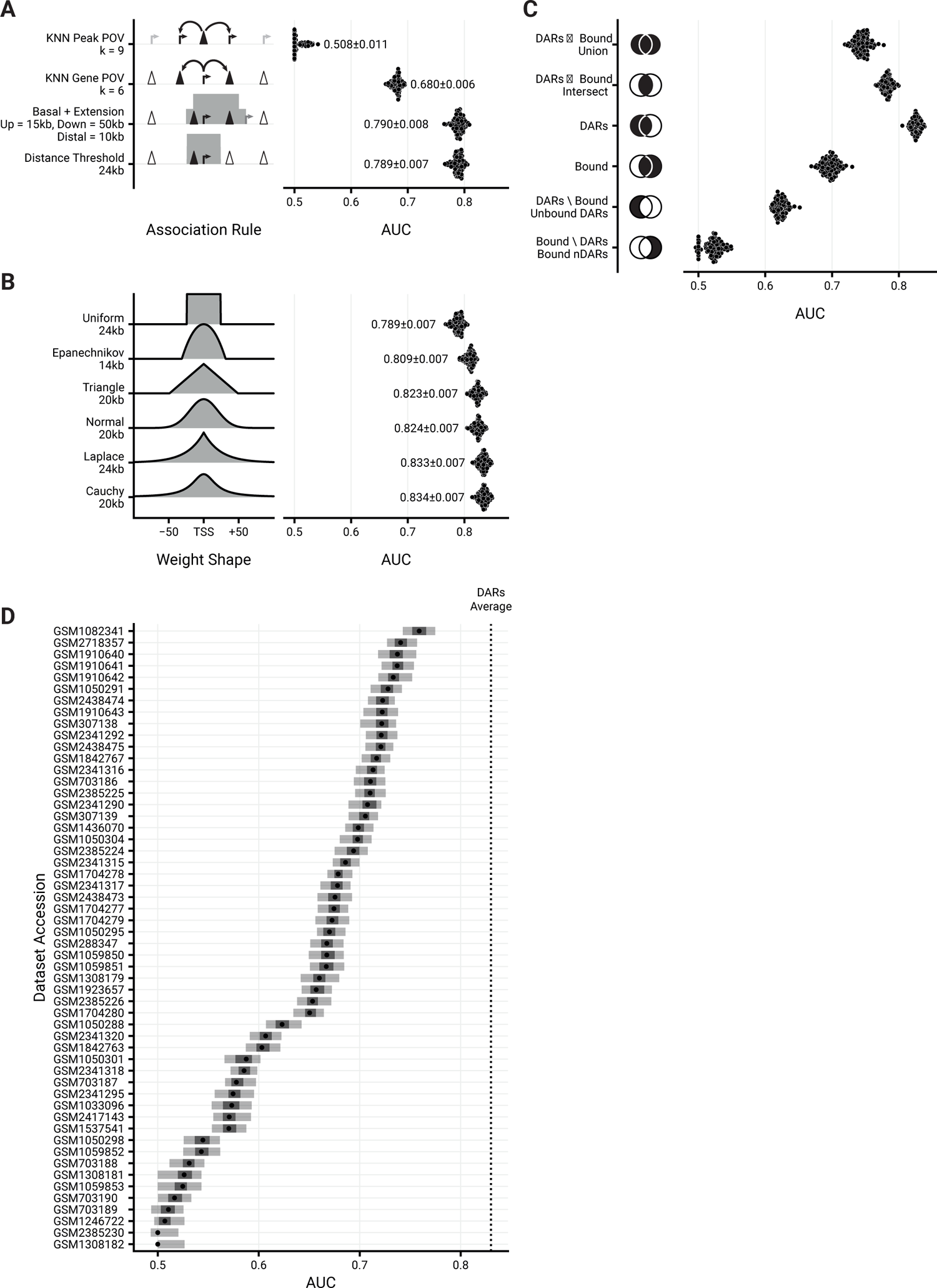
Characterization of different association rules and chromatin features for the prediction of transcriptional changes. **(A)** Predictive performance comparison of different association rules to discriminate downregulated DEGs upon SOX2 depletion at 2 hours from matched nDEGs, with sums of (n)DAR peaks as predictors. X-axis metric notes the area under the receiver operator characteristic curve. Dots represent 100 re-samplings of the matched nDEGs. Numbers represent mean ± standard deviation. Optimal parameters for every rule, such as distance = 25kb and k = 6, were chosen by performing a parameter sweep and choosing the parameter that minimized cross-validation error. **(B)** Like (A), but for various kernel functions instead of association rules, and weighted sums instead of plain sums as predictors. Left part gives visual indication of kernel shape. **(C)** Prediction of differential gene expression illustrate the predictive power of combinations of DARs and SOX2 binding measured by ChIPseq. Venn diagrams illustrate the set operations of SOX2 and ATACseq. **(D)** Predictive power of expression changes based on the many publicly available ChIPseq data for SOX2 (via Cistrome). The dotted line indicate the average predictive power of DARs.

**Figure S6.**
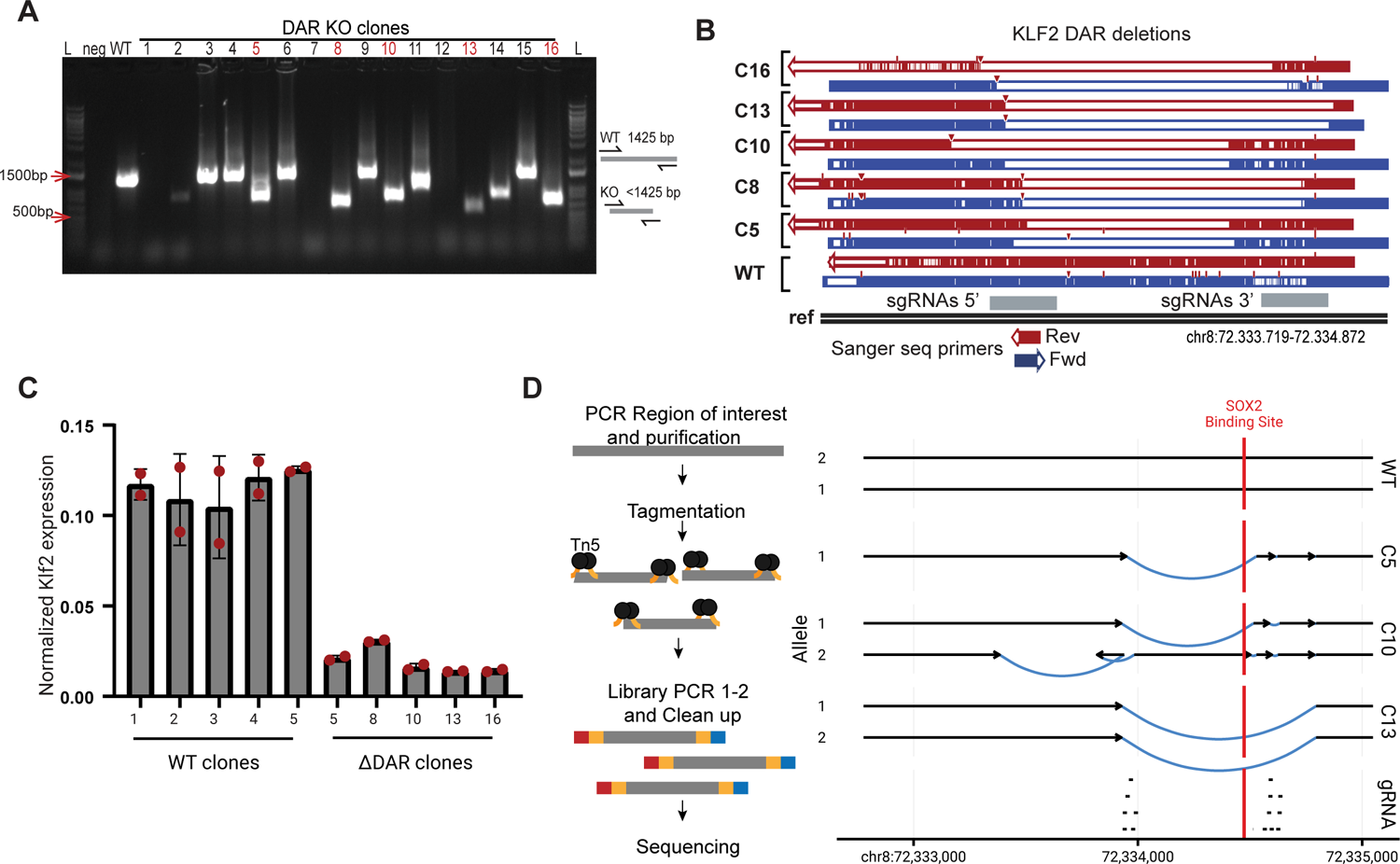
Validation of genome editing of DAR KO and *Klf2* gene expression. (**A**) Gel electrophoresis of PCR for genotyping disruption of the DAR region in clones from the gene edited SOX2-FKBP parental cell line. Primers amplifying the targeted regions were used to control for the homozygous disruption compared to WT amplification. L: ladder, Neg: water control, DAR KO clones: clones selected for genotyping. In red, clones selected for further experiments. (**B**) Validation of the disruption using Sanger sequencing in clones compared to non-edited clones. Blue: forward primer, red: reverse primer, grey: region targeted by sgRNAs 5’ and 3’ of the DAR. (**C**) RT-qPCR of *Klf2* expression, similar as Figure 5B, but using an alternative set of primers, in 5 parental clones and the DAR KO clones. Expression is relative to housekeeping gene *Rsp26*. (**D**) Left panel shows simplified overview of the amplicon sequencing procedure. Right panel shows the most likely assembly based on the amplicon sequencing of the DAR regions in WT and KO clones. Region targeted by sgRNAs are shown at the bottom. Arrows indicate the centromere to telomere orientation. Black lines shows assembled sequence that is not difference from the reference sequence. Blue lines show structural variants identified in the clones. Red indicates position of the SOX2 binding motif overlapping with a SOX2 ChIPseq peak.

