## Supplemental Figures for "Pioneer activity distinguishes activating from non-activating pluripotency transcription factor binding sites"

### Figure S1

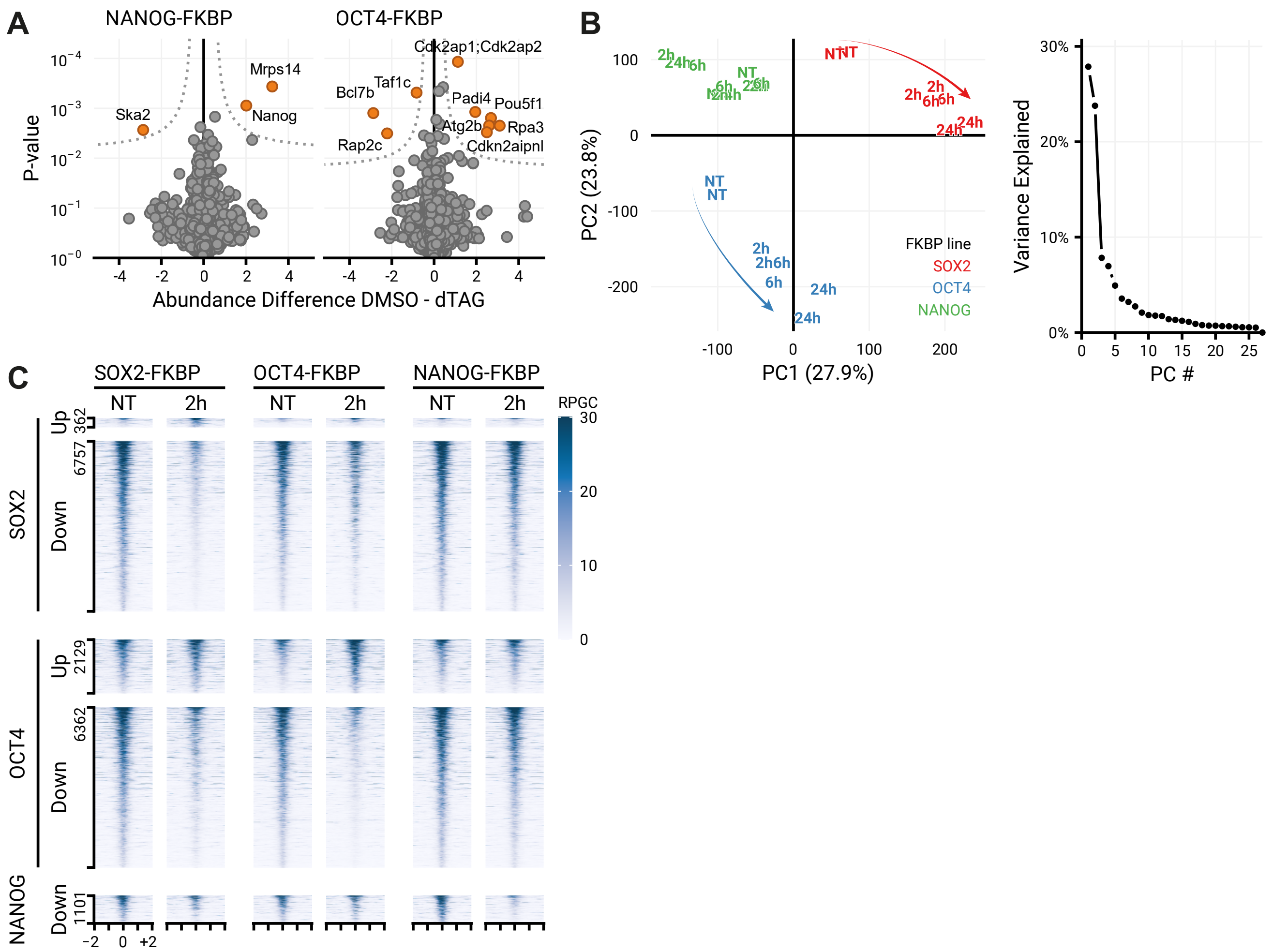

A
NANOG-FKBP
OCT4-FKBP
P-value
Abundance Difference DMSO - dTAG
Mrps14
Ska2
Nanog
Cdk2ap1;Cdk2ap2
Taf1c
Bcl7b
Padi4
Pou5f1
Atg2b
Rpa3
Rap2c
Cdkn2aipnl
B
PC1 (27.9%)
PC2 (23.8%)
FKBP line
SOX2
OCT4
NANOG
Variance Explained
PC #
C
SOX2-FKBP
OCT4-FKBP
NANOG-FKBP
NT
2h
RPGC
SOX2
OCT4
NANOG
Up
Down
362
6757
2129
6362
1101

**Figure S1 Quantification of changes the proteome and open chromatin landscape following depletion of pluripotency transcription factors**

**(A)** Quantitative mass spectrometry results showing the differential protein abundance comparing 30 minutes of dTAG-13 treated and untreated NANOG-FKBP and OCT4-FKBP cells. **(B)** Principal component analysis (PCA) scatter plot of accessibility data of the three FKBP-tagged lines indicating the first two principal components (PCs) (left). Arrows indicate major movements over time. On the right, the amount of variance explained by each principal component. **(C)** Tornado plot showing accessibility in the SOX2-, OCT4- and NANOG-FKBP-lines in untreated cells and following 2 hours of dTAG-13 treatment at differentially accessible regions for the indicated lines. Note that the 'Up' category in the NANOG-FKBP has been omitted for having too few peaks.

**Figure S2**

A

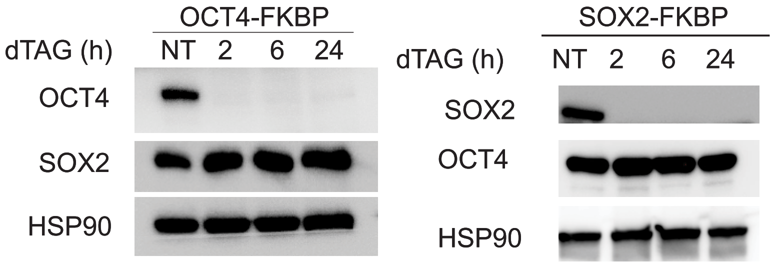
OCT4-FKBP
dTAG (h)
NT
2
6
24
OCT4
SOX2
HSP90
SOX2-FKBP
dTAG (h)
NT
2
6
24
SOX2
OCT4
HSP90

**Figure S2 OCT4 and SOX2 depletion does not affect protein levels of SOX2 and OCT4**
(A) Western blot showing the levels of OCT4 and SOX2 protein in the OCT4-FKBP cell line after 2, 6 and 24h of dTAG-13 treatment. HSP90 used as loading control. (B) The same as in (A) but in the SOX2-FKBP cell line.

### Figure S3

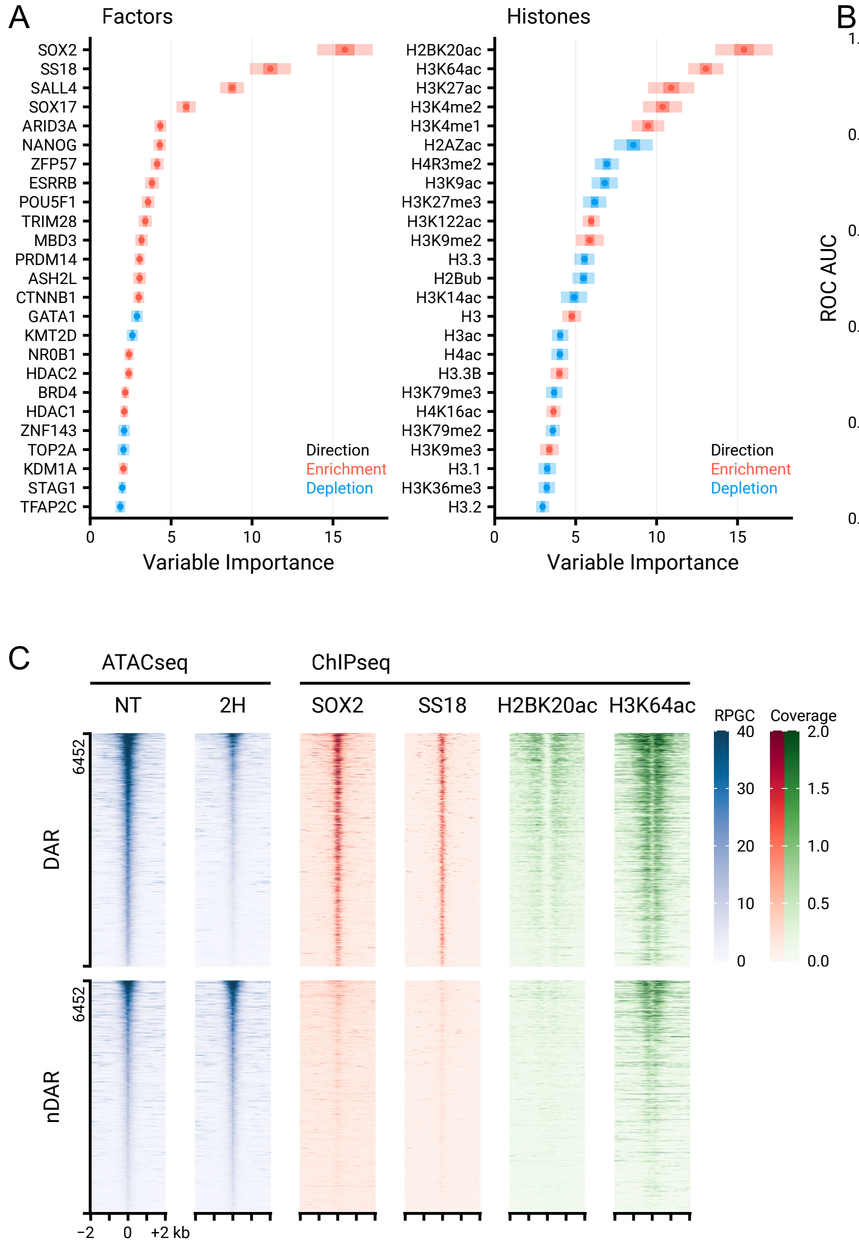

A
Factors
SOX2
SS18
SALL4
SOX17
ARID3A
NANOG
ZFP57
ESRRB
POU5F1
TRIM28
MBD3
PRDM14
ASH2L
CTNNB1
GATA1
KMT2D
NR0B1
HDAC2
BRD4
HDAC1
ZNF143
TOP2A
KDM1A
STAG1
TFAP2C
Direction
Enrichment
Depletion
Variable Importance
Histones
H2BK20ac
H3K64ac
H3K27ac
H3K4me2
H3K4me1
H2AZac
H4R3me2
H3K9ac
H3K27me3
H3K122ac
H3K9me2
H3.3
H2Bub
H3K14ac
H3
H3ac
H4ac
H3.3B
H3K79me3
H4K16ac
H3K79me2
H3K9me3
H3.1
H3K36me3
H3.2
Direction
Enrichment
Depletion
Variable Importance
B
ROC AUC
Cumulative number of datasets
Datasets
Factor
Histones
C
ATACseq
ChIPseq
NT
2H
SOX2
SS18
H2BK20ac
H3K64ac
RPGC
Coverage
6452
DAR
6452
nDAR
−2
0
+2 kb

**Figure S3 Random forest classification using Cistrome data reveals proteins and histone modifications that can predict differential accessibility following SOX2 depletion.**

**(A)** Left**,** top 25 (chromatin binding) factors in the Cistrome factors datasets whose overlap with all ATACseq peaks is predictive in random forest classification to discriminate the DAR from nDAR peaks. ATACseq peaks were extended by 300bp in both directions. Variable importance was calculated with subsampling inference, wherein the 95% confidence interval (CI) is indicated with a light colour, the 50% CI with a darker colour and the median with a point. Enrichment and depletion indicate higher average overlap in the DAR than nDAR categories respectively. Right, top 25 histone modifications using the Cistrome histone datasets.. **(B)** Area under the receiver operator characteristic curve indicating predictive performance using an increasing number of factors/histone modifications sorted by variable importance in discriminating DAR from nDAR peaks. Dotted lines indicate predictive performance of using all factors/histones for prediction (n=233/n=38). **(C)** Tornado plots showing ATACseq data for untreated and 2h dTAG-13 treated SOX2-FKBP cells and ChIPseq signal for the top two predictive datasets from the factors and histone databases over the DAR and nDAR peaks. RPGC = reads per genomic content, coverage indicates values in pre-processed data(Zheng et al., 2019).

### Figure S4

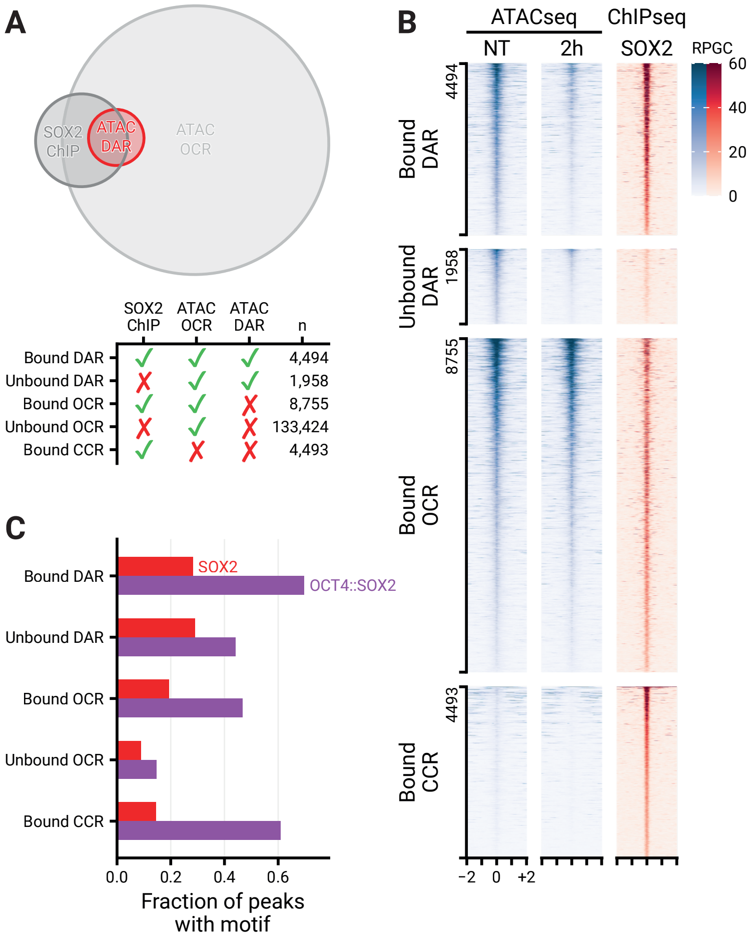
A
SOX2 ChIP
ATAC DAR
ATAC OCR
SOX2 ChIP
ATAC OCR
ATAC DAR
n
Bound DAR
Unbound DAR
Bound OCR
Unbound OCR
Bound CCR
4,494
1,958
8,755
133,424
4,493
B
ATACseq
ChIPseq
NT
2h
SOX2
RPGC
60
40
20
0
Bound DAR
4494
Unbound DAR
1958
Bound OCR
8755
Bound CCR
4493
−2
0
+2
C
SOX2
OCT4::SOX2
Bound DAR
Unbound DAR
Bound OCR
Unbound OCR
Bound CCR
0.0
0.2
0.4
0.6
Fraction of peaks with motif

**Figure S4 Quantitative and qualitative analysis of ATACseq and ChIPseq**

**(A)** Venn diagram showing the overlap between differentially accessible regions (DARs), all ATACseq peaks (OCR) and SOX2 DNA binding (SOX2 ChIP). Bottom panel shows the number of peaks in each overlap category. **(B)** Tornado plot showing accessibility before and after 2 hours of SOX2 depletion, along with SOX2 binding, for the sites in Figure 4A. **(C)** Fraction of peaks containing the Oct4::Sox2 or Sox2 DNA binding motifs, stratified by whether they are accessible peaks as measured by ATACseq, SOX2 binding sites as measured by ChIPseq, differentially accessible after 2 hours of SOX2 depletion or combinations thereof.

(OCR, Open chromatin regions. DAR, Differential accessible regions. CCR, closed chromatin regions)

### Figure S5

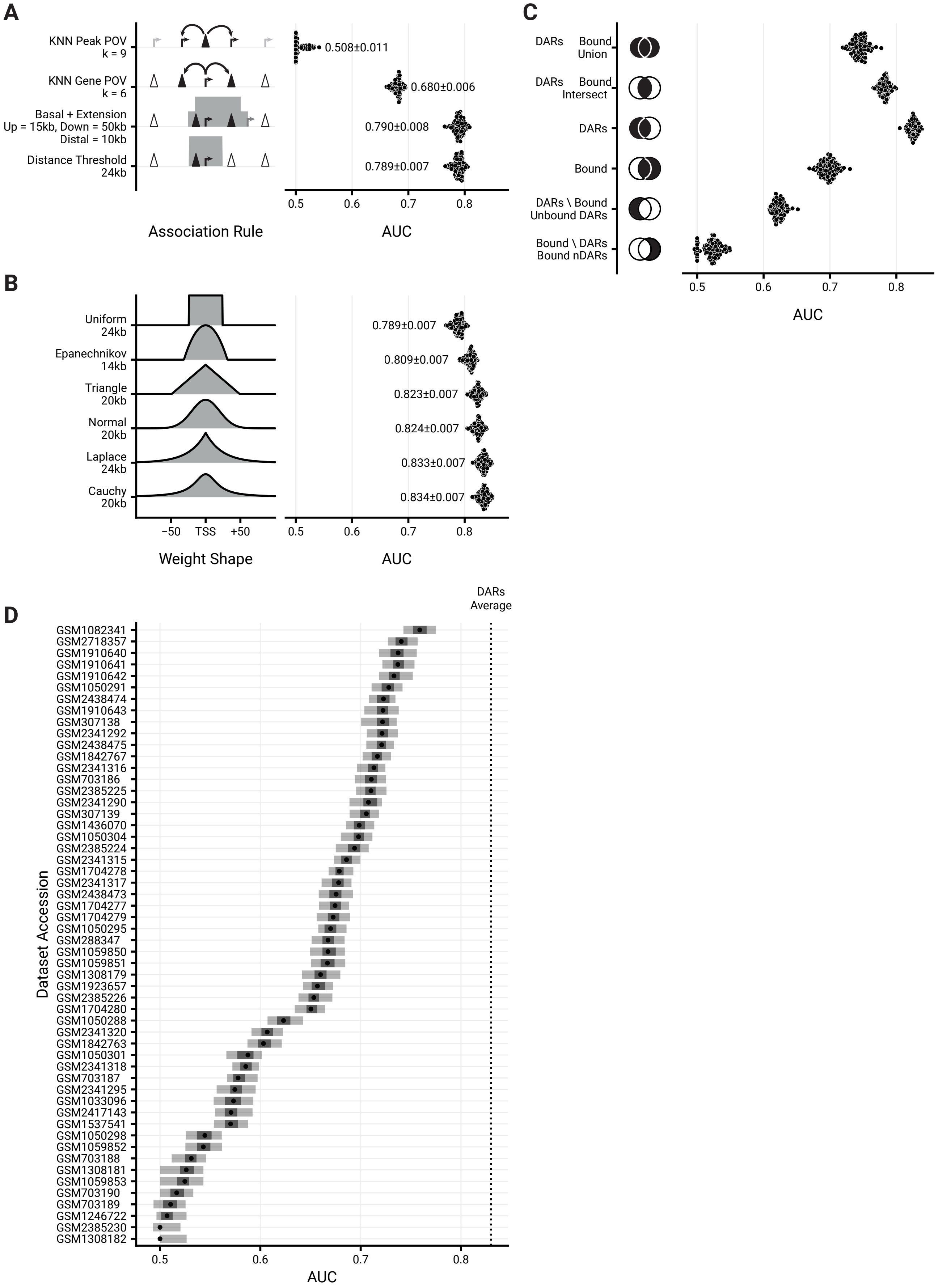
A
KNN Peak POV
k = 9
KNN Gene POV
k = 6
Basal + Extension
Up = 15kb, Down = 50kb
Distal = 10kb
Distance Threshold
24kb
0.508±0.011
0.680±0.006
0.790±0.008
0.789±0.007
0.5
0.6
0.7
0.8
Association Rule
AUC
B
Uniform
24kb
Epanechnikov
14kb
Triangle
20kb
Normal
20kb
Laplace
24kb
Cauchy
20kb
0.789±0.007
0.809±0.007
0.823±0.007
0.824±0.007
0.833±0.007
0.834±0.007
−50
TSS
+50
Weight Shape
AUC
C
DARs Bound
Union
DARs Bound
Intersect
DARs
Bound
DARs \ Bound
Unbound DARs
Bound \ DARs
Bound nDARs
AUC
D
DARs
Average
Dataset Accession
GSM1082341
GSM2718357
GSM1910640
GSM1910641
GSM1910642
GSM1050291
GSM2438474
GSM1910643
GSM307138
GSM2341292
GSM2438475
GSM1842767
GSM2341316
GSM703186
GSM2385225
GSM2341290
GSM307139
GSM1436070
GSM1050304
GSM2385224
GSM2341315
GSM1704278
GSM2341317
GSM2438473
GSM1704277
GSM1704279
GSM1050295
GSM288347
GSM1059850
GSM1059851
GSM1308179
GSM1923657
GSM2385226
GSM1704280
GSM1050288
GSM2341320
GSM1842763
GSM1050301
GSM2341318
GSM703187
GSM2341295
GSM1033096
GSM2417143
GSM1537541
GSM1050298
GSM1059852
GSM703188
GSM1308181
GSM1059853
GSM703190
GSM703189
GSM1246722
GSM2385230
GSM1308182
0.5
0.6
0.7
0.8
AUC

**Figure S5 Characterization of different association rules and chromatin features for the prediction of transcriptional changes.**

**(A)** Predictive performance comparison of different association rules to discriminate downregulated DEGs upon SOX2 depletion at 2 hours from matched nDEGs, with sums of (n)DAR peaks as predictors. X-axis metric notes the area under the receiver operator characteristic curve. Dots represent 100 re-samplings of the matched nDEGs. Numbers represent mean ± standard deviation. Optimal parameters for every rule, such as distance = 25kb and k = 6, were chosen by performing a parameter sweep and choosing the parameter that minimized cross-validation error. **(B)** Like (A), but for various kernel functions instead of association rules, and weighted sums instead of plain sums as predictors. Left part gives visual indication of kernel shape. **(C)** Prediction of differential gene expression illustrate the predictive power of combinations of DARs and SOX2 binding measured by ChIPseq. Venn diagrams illustrate the set operations of SOX2 and ATACseq. **(D)** Predictive power of expression changes based on the many publicly available ChIPseq data for SOX2 (via Cistrome). The dotted line indicate the average predictive power of DARs.

### Figure S6

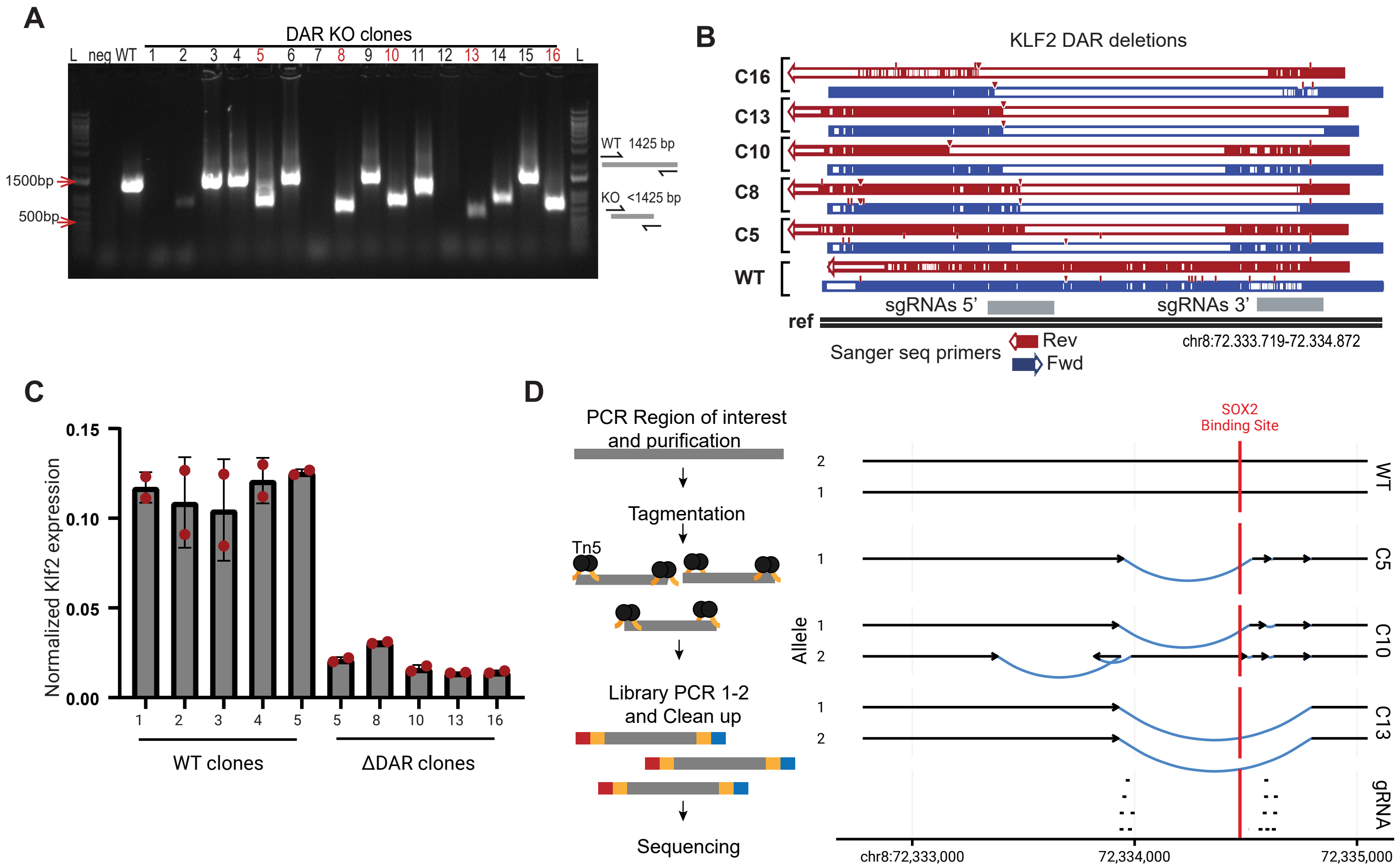
A
DAR KO clones
L neg WT 1 2 3 4 5 6 7 8 9 10 11 12 13 14 15 16 L
1500bp
500bp
WT 1425 bp
KO <1425 bp
B
KLF2 DAR deletions
C16
C13
C10
C8
C5
WT
sgRNAs 5'
sgRNAs 3'
ref
Sanger seq primers
Rev
Fwd
chr8:72.333.719-72.334.872
C
Normalized Klf2 expression
0.15
0.10
0.05
0.00
1 2 3 4 5 5 8 10 13 16
WT clones
ΔDAR clones
D
PCR Region of interest and purification
Tagmentation
Tn5
Library PCR 1-2 and Clean up
Sequencing
SOX2 Binding Site
Allele
WT
C5
C10
C13
gRNA
chr8:72,333,000
72,334,000
72,335,000

**Figure S6 Validation of genome editing of DAR KO and *Klf2* gene expression**

(**A**) Gel electrophoresis of PCR for genotyping disruption of the DAR region in clones from the gene edited SOX2-FKBP parental cell line. Primers amplifying the targeted regions were used to control for the homozygous disruption compared to WT amplification. L: ladder, Neg: water control, DAR KO clones: clones selected for genotyping. In red, clones selected for further experiments. (**B**) Validation of the disruption using Sanger sequencing in clones compared to non-edited clones. Blue: forward primer, red: reverse primer, grey: region targeted by sgRNAs 5' and 3' of the DAR. (**C**) RT-qPCR of *Klf2* expression, similar as Figure 5B, but using an alternative set of primers, in 5 parental clones and the DAR KO clones. Expression is relative to housekeeping gene *Rsp26*. (**D**) Left panel shows simplified overview of the amplicon sequencing procedure. Right panel shows the most likely assembly based on the amplicon sequencing of the DAR regions in WT and KO clones. Region targeted by sgRNAs are shown at the bottom. Arrows indicate the centromere to telomere orientation. Black lines shows assembled sequence that is not difference from the reference sequence. Blue lines show structural variants identified in the clones. Red indicates position of the SOX2 binding motif overlapping with a SOX2 ChIPseq peak.
